# Complementary benefits of multivariate and hierarchical models for identifying individual differences in cognitive control

**DOI:** 10.1101/2024.04.24.591032

**Authors:** Michael C. Freund, Ruiqi Chen, Gang Chen, Todd S. Braver

## Abstract

Understanding individual differences in cognitive control is a central goal in psychology and neuroscience. Reliably measuring these differences, however, has proven extremely challenging, at least when using standard measures in cognitive neuroscience such as response times or task-based fMRI activity. While prior work has pinpointed the source of the issue — the vast amount of cross-trial variability within these measures — solutions remain elusive. Here, we propose one potential way forward: an analytic framework that combines hierarchical Bayesian modeling with multivariate decoding of trial-level fMRI data. Using this framework and longitudinal data from the Dual Mechanisms of Cognitive Control project, we estimated individuals’ neural responses associated with cognitive control within a color-word Stroop task, then assessed the reliability of these individuals’ responses across a time interval of several months. We show that in many prefrontal and parietal brain regions, test–retest reliability was near maximal, and that only hierarchical models were able to reveal this state of affairs. Further, when compared to traditional univariate contrasts, multivariate decoding enabled individual-level correlations to be estimated with significantly greater precision. We specifically link these improvements in precision to the optimized suppression of cross-trial variability in decoding. Together, these findings not only indicate that cognitive control-related neural responses individuate people in a highly stable manner across time, but also suggest that integrating hierarchical and multivariate models provides a powerful approach for investigating individual differences in cognitive control, one that can effectively address the issue of high-variability measures.

## 1 Introduction

A major goal within psychology and neuroscience is to understand the mechanisms that give rise to psychological diversity. How are two minds alike or distinct, in terms of psychological processes? What neural mechanisms underlie this variability? Such questions of *individual differences* are incredibly important to study: not only do they intrinsically interest many, but they also provide a means to test virtually any cognitive psychological theory (*cf.*, Underwood, 1975), as well as yield direct clinical and educational applications (Cole et al., 2014; Diamond, 2013; Engle, 2002). Spurred by these motivations, cognitive neuroscientists have often sought to identify measures of human brain activity that can be used as markers of individual differences. Yet, this enterprise has proven highly difficult, at least when using common non-invasive measures of brain activity, such as task-based fMRI.

One of the main challenges in studying individual differences can be traced to issues of measurement. In the laboratory, people may score differently on experimental tasks due to different reasons, many of which have nothing to do with cognitive properties of the individual that are stable over time, here termed *cognitive traits*, but instead with properties of the particular measurement occasion, here termed nuisance or *noise* factors. Thus, when reasoning about cognitive traits, researchers run the risk of misattributing a particular finding or pattern of results to cognitive traits that in fact arise from more transient (i.e., noise) factors.

The statistic of *test-retest reliability* is instrumental in mitigating this risk. In particular, test–retest reliability is estimated by repeatedly acquiring the same measures from the same sample of individuals over an extended period of time (typically on the order of days, at minimum). In such repeated-measures designs, differences observed between individuals that are constant over *test repetitions* (also known as “sessions”) are assumed to reflect traits, while changes within individuals over repetitions are assumed to reflect noise (which may include systematic longitudinal effects, such as learning). The relative proportion of trait variance defines the test-retest reliability, or the “traitness” of the measure in question. Yet, despite this grounding theoretical framework, in practice it has not been easy to identify strong individual differences in cognition (conventionally, r > 0.7; Matheson, 2019). This difficulty has been particularly salient with measures derived from classical cognitive experimental tasks (Hedge et al., 2018) and adaptations of these paradigms for fMRI. Perhaps most confoundingly, however, poor test-retest reliability has often been reported in psychological domains typically assumed to be subject to strong individual differences, for example cognitive control and working memory (Elliott et al., 2020). Such failures have invited pessimism over the usefulness of these popular methods for individual differences research (Elliott et al., 2020).

Much of these pessimistic conclusions, however, overlook a nuanced theoretical issue concerning the structure of noise variability: it is hierarchical, and using methods that cannot account for this hierarchical structure will lead to an overly pessimistic estimate of reliability (Chen et al., 2021). To elaborate, within the standard test-retest design, noise variability can be decomposed into (at least) two levels: trial-level variability (i.e., changes over trials, within repetition) and repetition-level variability (i.e., changes over repetitions, aggregated over trials). Overwhelmingly, prior work has estimated reliability through a two-stage summary statistic approach, which involves first aggregating a measure over trials, then assessing the relative proportion of trait variability via linear correlation (see Figure 1 for a walkthrough; see also Haines et al., 2020). But, because the estimation of the linear correlation does not use any information regarding the amount of trial-level variability, the summary statistic approach essentially “conceals” the negative impact such variability has on estimated test–retest reliability. In other words, summary statistic estimates of test–retest reliability are underestimated (Figure 1). The amount of underestimation depends in part on how strongly the measure differs across trials, and how effective the first-step aggregation was at reducing the standard errors of the scores. Theoretically, it is possible that the bias is small, but true physiological and behavioral measures vary too strongly over trials for this approach to be accurate in reality (i.e., given attainable amounts of data; Rouder et al., 2023).

**Figure 1:**
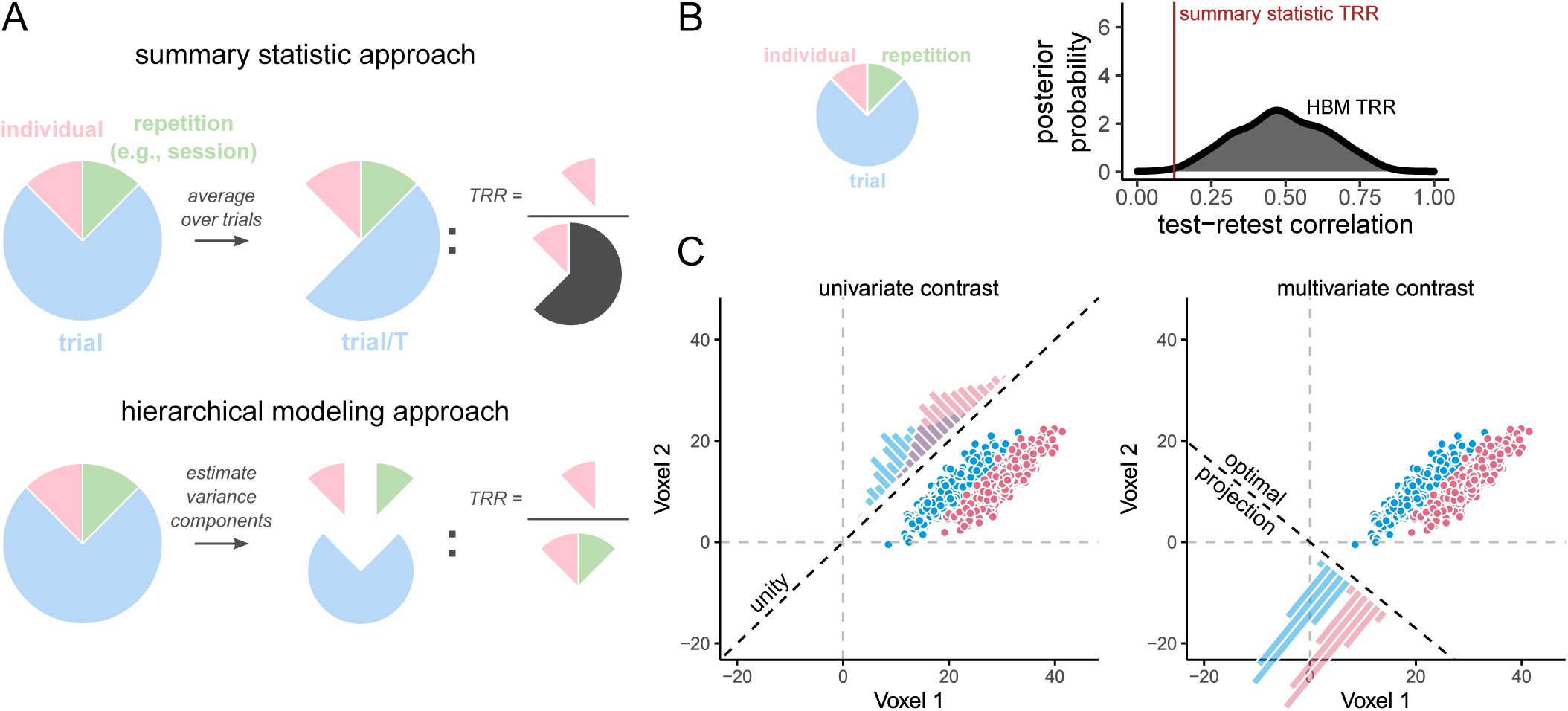
Graphical intuition for test-retest reliability and benefits of multivariate pattern analysis. **A, B**, Two approaches to estimating test-retest reliability: summary statistic versus hierarchical modeling. Psychological experimental tasks are typically composed of hierarchically structured “levels”: trials are completed within testing repetitions (e.g., sessions) within individuals. Measures derived from such tasks exhibit some amount of variability at each of these levels: for example, a person’s performance may fluctuate over trials, across different days (repetitions), while also reliably differing over time from another person’s average performance (individual differences). Rigorously studying individual differences requires disentangling individual variability from variability at the other levels. The extent to which a measure supports such disentangling is quantified by test–restest reliability (TRR). **A, top**, The summary statistic approach estimates reliability by first aggregating over trials per individual, attempting to quash trial-level variability prior to computing the proportion of individual-level variance. Despite this aggregation, however, a potentially substantial amount of residual trial-level variability remains “concealed” in the aggregated scores, and thereby in the denominator of the reliability statistic. **A, bottom**, In hierarchical approaches, however, the variance is decomposed into separate components, and trial-level variability is explicitly factored out of the test–retest reliability computation. **B**, When trial-level variability is high (pie chart), the test–retest reliability estimated through summary statistical approaches (red vertical line) will tend to be shrunken relative to estimates from hierarchical Bayesian models (HBM; the central tendency of the grey density indicates the average posterior test–retest correlation). By contrast, in hierarchical models, such heightened trial-level variability would instead manifest as decreased estimation precision (spread of grey density). Unfortunately, both of these scenarios would impair inferential power. **D**, Common forms of multivariate pattern analysis attenuate trial-level variability. Left and right panels show the same (simulated) fMRI data, acquired from a hypothetical region of interest composed of two voxels (x and y axes). Individual trials of a Stroop task evoked particular patterns of activity across these two voxels (points). Each trial belongs to one of two conditions, *incongruent* (red) or *congruent* (blue), whose difference forms the Stroop effect contrast. Both univariate analysis and multivariate pattern analysis can be used to compute the Stroop effect contrast. **Left panel**, In a univariate analysis, measures from different voxels are aggregated together *uniformly*, by taking the spatial mean (i.e., each voxel is weighted equally then summed). Graphically, this corresponds to projecting the condition means onto the unity line, which runs through the origin and *x* = 1, *y* = 1 (diagonal dotted line). Yet, depending on the shape and relative configuration of these distributions, such a projection may lead to a highly variable measure (overlapping histograms along unity line). **Right panel**, In a multivariate pattern analysis, measures from different voxels can be aggregated together *optimally*. Graphically, this corresponds to projecting the condition means onto the line that yields the most separation between projected classes (i.e., minimizing overlap between the histograms of the projections). As a result, this procedure can lead to projections in which trial-level variance is squashed relative to univariate analysis.

This consideration has motivated an alternative, hierarchical Bayesian modeling approach for assessing individual differences (Chen et al., 2021). A hierarchical approach does not attempt to squash trial-level variability through aggregation, but instead, it attempts to directly estimate the magnitude of each component of variability from the disaggregated trial-level measures. In this way, the negative impact of trial-level noise on reliability can be explicitly factored out (Figure 1B). In line with this reasoning, results from studies adopting a hierarchical Bayesian approach to individual differences suggest a more optimistic outlook, in which reliable individual differences can in some cases be identified after accounting for the contributions of trial-level noise (Chen et al., 2021; Haines et al., 2020; Snijder et al., 2023).

Nevertheless, these studies have also illustrated that the hierarchical Bayesian approach is not a panacea for investigating individual differences with high-variability measures (Rouder et al., 2023). This is because the issues caused by high trial-level variability are not fully circumvented in hierarchical Bayesian models. Instead, the issues are redirected, so that they manifest differently within the model estimates. That is, instead of diminishing reliability, increasing trial-level variability leads to *more uncertainty* in the reliability estimate (Figure 1C). Within a recent report, for example, hierarchical Bayesian reliability estimates of fMRI contrasts within select regions of interest were typically higher than their summary statistic counterparts, but were also highly uncertain, with the lower-bound confidence interval generally located well below zero (Chen et al., 2021). This “redirection” is invaluable, in that it clarifies a major source of the problems researchers have face when trying to measure individual differences: the overwhelming amount of trial-level variability leads to less certainty at the individual level. However, it does not solve the problem itself.

A plausible remedy for these issues is provided by multivariate pattern analysis (MVPA) decoding (Haxby et al., 2014; Hebart & Baker, 2018; Naselaris et al., 2011). By exploiting the high spatial dimensionality of fMRI, MVPA decoding can suppress trial-level variability by identifying dimensions of neural coding that are relatively insensitive to such fluctuations (Figure 1D). Yet generally, prior work on individual differences has instead opted to use more traditional univariate models, which can be much more vulnerable to trial-level variability (Elliott et al., 2020). This combination of facts suggests a constructive hypothesis: MVPA decoding can improve the power of individual differences analyses by increasing the certainty (i.e., precision) of the estimated trait-level parameters. Some indirect evidence is indeed in line with this hypothesis (Kragel et al., 2021; Xu et al., 2018; Yoo et al., 2019), however, a direct and more extensive investigation of these issues is warranted, in particular within the domain of task-related fMRI (*cf.*, resting state).

Here, using a subset of densely-sampled individuals (N = 27, 6 sessions each) acquired as part of the Dual Mechanisms of Cognitive Control fMRI dataset (Braver et al., 2021), we compared the sensitivity of various approaches to individual differences analyses using a classical fMRI measure of cognitive control function, the Stroop-effect contrast (incongruent versus congruent trials), as derived from the color-word Stroop, 1935 task. We compared the reliability of a traditional univariate version of this measure to a multivariate contrast, obtained from MVPA decoding, that we specifically constructed to suppress trial-level variability. To provide an accurate and complete comparison, we performed these analyses within the context of a hierarchical Bayesian modeling framework for individual differences (Chen et al., 2021) and subsequently compared the results to those derived from a summary statistic approach. In line with theory, we found that relative to univariate models, MVPA decoding improved the ability to identify strong individual differences in fMRI measures of cognitive control from frontoparietal cortex. In a hierarchical Bayesian framework, these improvements manifested as increased estimation certainty (i.e., precision) of individual difference parameters. These results highlight the efficacy of a joint hierarchical Bayesian– multivariate decoding approach, in which the statistical accuracy of hierarchical Bayesian models is boosted by the statistical power of MVPA decoding, and suggest that non-invasive neural measures like task-based fMRI may be an adequate means for addressing questions regarding individual differences in cognition.

## 2 Method

### 2.1 Mathematical Notation

The transpose operation is written ^⊺^. Non-bold font is used for scalars while lowercase bold is used for column vectors and uppercase bold for matrices. Italicized subscripts are used to index sets of scalars, vectors, or matrices: for example, **X***_i,j_* for *i* ∈ 1*, …, I* and *j* ∈ 1*, …, J* refers to the *i,j*-th matrix in a set of *IJ* matrices. Importantly, this subscript notation should not be confused with indices for elements of individual vectors or matrices. To avoid such confusion, we write element-wise indices with parenthetical superscripts: for example, 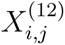 refers to the element in the first row and second column of matrix **X***_i,j_*. For clarity within inline text, index notation will typically be omitted.

The specific notation used for factor indices are listed in Table 2.1.

**Table 1:**
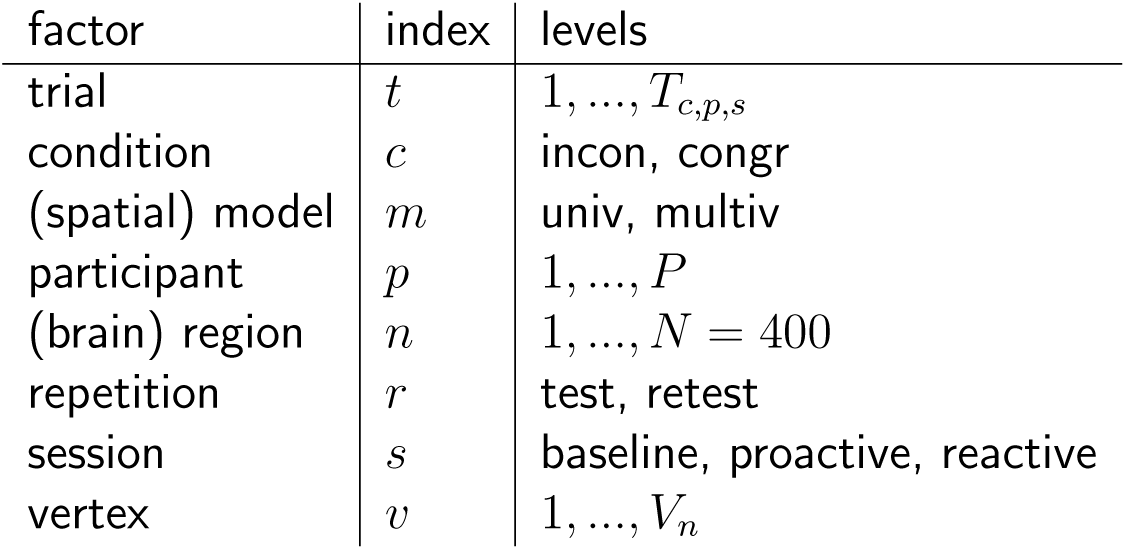
Experimental factors and indices.

| factor | index | levels |
| --- | --- | --- |
| trial | $t$ | $1, \dots, T_{c,p,s}$ |
| condition | $c$ | incon, congr |
| (spatial) model | $m$ | univ, multiv |
| participant | $p$ | $1, \dots, P$ |
| (brain) region | $n$ | $1, \dots, N = 400$ |
| repetition | $r$ | test, retest |
| session | $s$ | baseline, proactive, reactive |
| vertex | $v$ | $1, \dots, V_n$ |

**Table 2:** Regions of interest and associated test–retest reliability statistics. Displayed are key statistics from Schaefer-atlas parcels that exhibited the strongest test–retest reliability in the multivariate Stroop activation contrast. Asterisks by parcel names indicate regions of interest associated with cognitive control demands, defined in a prior report (Braver et al., 2021; 11 out of 35 also exhibit strong test–retest reliability). Per region, the strength of population-level univariate activation and multivariate discriminability are summarized in the *t*^+^ statistics. Test–retest reliability in univariate activation contrasts (Univariate TRR) and in multivariate contrasts (Multivariate TRR) are also displayed. The maximum *a posteriori* estimate (MAP) indicates the central tendency of posterior TRR correlations (i.e., the most likely point estimate), while the 5^th^ percentile of the posterior (5%ile) indicates a lower-bound on the estimate (i.e., the uncertainty in the estimate). Likewise, the 95^th^ percentile indicates an upper bound. Rows are sorted by the lower bound. The Variability Ratio indicates the trial/subject ratio (see Method; here, this ratio is not log transformed). Intra-class correlation coefficient estimates (ICC) are also displayed.

| Parcel (Schaefer 400-17) | Univ. $t^+$ | Multiv. $t^+$ | Univariate TRR | | | | | | Multivariate TRR | | | | | |
| --- | --- | --- | --- | --- | --- | --- | --- | --- | --- | --- | --- | --- | --- | --- |
|  |  |  | ICC | MAP | Mean | 5%ile | 95%ile | VR | ICC | MAP | Mean | 5%ile | 95%ile | VR |
| LH DorsAttnA SPL 3 | 4.58 | 3.91 | 0.61 | 0.97 | 0.88 | 0.46 | 0.99 | 6.12 | 0.87 | 0.94 | 0.92 | 0.75 | 0.98 | 2.24 |
| RH DorsAttnA SPL 4 | 2.23 | 2.56 | 0.79 | 0.99 | 0.94 | 0.67 | 0.99 | 4.80 | 0.75 | 0.97 | 0.94 | 0.74 | 0.99 | 3.43 |
| LH ContA IPS 5* | 4.65 | 5.87 | 0.54 | 0.98 | 0.90 | 0.46 | 0.99 | 7.01 | 0.71 | 0.99 | 0.93 | 0.67 | 0.99 | 4.31 |
| LH VisCent ExStr 11 | 1.82 | 2.63 | 0.35 | 0.68 | 0.56 | -0.16 | 0.92 | 7.90 | 0.67 | 0.95 | 0.91 | 0.63 | 0.99 | 4.05 |
| LH DorsAttnA SPL 6 | 0.60 | 3.04 | 0.36 | 0.95 | 0.63 | -0.51 | 0.97 | 12.39 | 0.78 | 0.98 | 0.92 | 0.62 | 0.99 | 4.69 |
| RH DorsAttnA SPL 7 | 0.43 | 1.70 | 0.47 | 0.95 | 0.81 | 0.23 | 0.98 | 7.64 | 0.66 | 0.96 | 0.90 | 0.59 | 0.99 | 4.37 |
| LH DorsAttnA SPL 7 | 1.20 | 3.51 | 0.14 | 0.68 | 0.35 | -0.66 | 0.92 | 15.68 | 0.67 | 0.99 | 0.92 | 0.59 | 0.99 | 4.93 |
| LH DorsAttnB PrCv 1* | 4.02 | 5.56 | 0.59 | 0.98 | 0.90 | 0.55 | 0.99 | 5.06 | 0.65 | 0.87 | 0.85 | 0.55 | 0.97 | 3.40 |
| LH DorsAttnA SPL 1 | 2.87 | 4.03 | 0.37 | 0.97 | 0.85 | 0.33 | 0.99 | 6.85 | 0.70 | 0.86 | 0.79 | 0.52 | 0.93 | 2.92 |
| LH DorsAttnA SPL 2 | 2.02 | 3.63 | 0.34 | 0.78 | 0.56 | -0.41 | 0.95 | 10.38 | 0.71 | 0.87 | 0.82 | 0.51 | 0.95 | 3.42 |
| RH DorsAttnA SPL 5 | -0.31 | 1.79 | 0.48 | 0.77 | 0.59 | -0.18 | 0.94 | 7.19 | 0.65 | 0.99 | 0.91 | 0.51 | 0.99 | 6.61 |
| RH DorsAttnA SPL 2 | 2.91 | 4.22 | 0.58 | 0.97 | 0.89 | 0.52 | 0.99 | 5.49 | 0.67 | 0.88 | 0.83 | 0.50 | 0.97 | 3.76 |
| LH DorsAttnA SPL 4* | 4.76 | 5.25 | 0.71 | 0.99 | 0.95 | 0.76 | 1.00 | 4.17 | 0.58 | 0.85 | 0.80 | 0.49 | 0.95 | 3.14 |
| RH ContA IPS 1 | 2.81 | 4.72 | 0.42 | 0.91 | 0.67 | -0.26 | 0.97 | 11.37 | 0.64 | 0.87 | 0.83 | 0.49 | 0.97 | 4.09 |
| LH DorsAttnA ParOcc 1 | -2.69 | 4.04 | -0.24 | -0.86 | -0.19 | -0.93 | 0.84 | 30.25 | 0.58 | 0.98 | 0.88 | 0.48 | 0.99 | 5.76 |
| LH DefaultB IPL 1 | -2.22 | 3.45 | -0.11 | 0.49 | 0.23 | -0.74 | 0.91 | 11.66 | 0.65 | 0.93 | 0.85 | 0.47 | 0.98 | 5.14 |
| LH DefaultB PFCi 2* | 4.76 | 5.44 | 0.29 | 0.88 | 0.52 | -0.55 | 0.96 | 12.74 | 0.58 | 0.98 | 0.87 | 0.47 | 0.99 | 5.55 |
| LH ContA IPS 4* | 4.94 | 5.37 | 0.62 | 0.98 | 0.90 | 0.49 | 0.99 | 7.26 | 0.64 | 0.91 | 0.84 | 0.46 | 0.97 | 4.58 |
| RH ContA PFCi 3* | 3.80 | 4.23 | 0.36 | 0.69 | 0.45 | -0.57 | 0.94 | 12.37 | 0.53 | 0.98 | 0.89 | 0.46 | 0.99 | 6.30 |
| LH ContA Temp 1 | -0.69 | 4.31 | 0.21 | 0.94 | 0.63 | -0.41 | 0.97 | 12.06 | 0.60 | 0.87 | 0.82 | 0.46 | 0.96 | 3.98 |
| LH DorsAttnA TempOcc 4 | 4.06 | 4.70 | 0.26 | 0.92 | 0.38 | -0.76 | 0.94 | 19.27 | 0.62 | 0.82 | 0.81 | 0.45 | 0.96 | 4.43 |
| LH DorsAttnA ParOcc 2 | 1.12 | 5.33 | 0.59 | 0.99 | 0.93 | 0.61 | 1.00 | 6.03 | 0.48 | 0.96 | 0.88 | 0.44 | 0.99 | 5.95 |
| RH DefaultB Temp 2 | 3.04 | 2.68 | 0.07 | 0.07 | 0.01 | -0.76 | 0.77 | 10.12 | 0.50 | 0.93 | 0.84 | 0.43 | 0.98 | 5.37 |
| RH DorsAttnA TempOcc 2 | 2.11 | 2.64 | 0.49 | 0.98 | 0.82 | 0.08 | 0.99 | 8.27 | 0.48 | 0.98 | 0.87 | 0.42 | 0.99 | 5.97 |
| LH VisCent ExStr 10 | -1.61 | 2.56 | 0.15 | 0.74 | 0.09 | -0.88 | 0.91 | 24.11 | 0.26 | 0.98 | 0.88 | 0.41 | 0.99 | 6.79 |
| LH DorsAttnB PostC 5 | -0.03 | 2.21 | 0.51 | 0.95 | 0.77 | -0.08 | 0.98 | 8.35 | 0.47 | 0.95 | 0.85 | 0.41 | 0.98 | 5.65 |
| LH SalVentAttnA ParOper 1 | -3.18 | 3.92 | 0.16 | 0.85 | 0.35 | -0.67 | 0.93 | 12.44 | 0.60 | 0.95 | 0.85 | 0.40 | 0.98 | 6.16 |
| RH ContA IPS 4 | 2.67 | 3.91 | 0.62 | 0.94 | 0.72 | -0.17 | 0.97 | 10.42 | 0.50 | 0.84 | 0.83 | 0.40 | 0.98 | 5.28 |
| LH DorsAttnB PostC 3 | 0.75 | 4.20 | 0.57 | 0.97 | 0.86 | 0.28 | 0.99 | 8.48 | 0.61 | 0.84 | 0.77 | 0.39 | 0.94 | 4.16 |
| LH ContA PFCd 1 | -2.26 | 2.67 | 0.08 | 0.88 | 0.42 | -0.74 | 0.95 | 19.45 | 0.53 | 0.93 | 0.83 | 0.34 | 0.98 | 5.89 |
| RH SalVentAttnA ParOper 1 | -3.35 | 4.00 | 0.03 | 0.56 | 0.22 | -0.78 | 0.92 | 13.59 | 0.55 | 0.94 | 0.83 | 0.34 | 0.98 | 6.46 |
| RH DefaultC IPL 1 | -3.38 | 3.75 | 0.33 | 0.83 | 0.61 | -0.31 | 0.96 | 10.74 | 0.45 | 0.96 | 0.82 | 0.33 | 0.98 | 6.84 |
| LH DorsAttnB FEF 2* | 1.22 | 3.46 | 0.32 | 0.92 | 0.48 | -0.71 | 0.96 | 21.02 | 0.46 | 0.93 | 0.84 | 0.33 | 0.98 | 6.87 |
| RH ContB IPL 4* | 4.34 | 4.03 | 0.55 | 0.91 | 0.74 | -0.18 | 0.98 | 11.29 | 0.46 | 0.86 | 0.78 | 0.31 | 0.97 | 5.18 |
| RH DorsAttnA SPL 8 | -2.83 | 1.42 | -0.07 | -0.63 | -0.11 | -0.89 | 0.86 | 18.76 | 0.50 | 0.97 | 0.83 | 0.29 | 0.98 | 7.48 |
| LH DorsAttnA SPL 5 | 1.77 | 3.64 | 0.37 | 0.95 | 0.74 | 0.06 | 0.97 | 8.22 | 0.48 | 0.95 | 0.82 | 0.28 | 0.98 | 6.09 |
| RH ContC pCun 1 | -2.64 | 1.53 | 0.63 | 0.98 | 0.90 | 0.48 | 0.99 | 6.70 | 0.51 | 0.98 | 0.84 | 0.27 | 0.99 | 7.92 |
| LH ContA IPS 3* | 5.95 | 7.11 | 0.40 | 0.93 | 0.80 | 0.18 | 0.98 | 7.77 | 0.54 | 0.68 | 0.64 | 0.26 | 0.87 | 3.83 |
| LH ContA PFCi 1* | 4.33 | 6.82 | 0.56 | 0.98 | 0.86 | 0.37 | 0.99 | 7.19 | 0.43 | 0.80 | 0.76 | 0.25 | 0.96 | 5.33 |
| LH ContA IPS 1* | 3.41 | 5.79 | 0.59 | 0.99 | 0.92 | 0.57 | 0.99 | 5.61 | 0.53 | 0.69 | 0.65 | 0.25 | 0.88 | 3.93 |

### 2.2 Subjects

Data were obtained as part of the Dual Mechanisms of Cognitive Control project (Braver et al., 2021). All subjects consented to participate in the study under Washington University IRB protocols. At the time of analysis, 32 subjects had completed the test–retest component of this project. We selected 27 of these subjects (number of females = 16, age range at initial session = [19, 42]) to include in the present analyses. This selection ensured that all subjects in our sample had complete data for all sessions that met minimal quality-control criteria (for each scanning run, complete behavioral and fMRI measures, low numbers of missed responses, and motion levels and dropout artifacts that were qualitatively judged to be modest). Of these subjects, 15 had also participated in the Human Connectome Project Young Adult study (Van Essen et al., 2013).

### 2.3 Design

Much of the debate over test-retest reliability has focused on the color-word Stroop, 1935 task as a paradigmatic example (e.g., Haines et al., 2020; Hedge et al., 2018; Rouder and Haaf, 2019). In order to directly address this debate, we focused exclusively on data from the Stroop task within the larger DMCC project dataset (Braver et al., 2021). In the Stroop task, names of colors were visually displayed in various nameable hues, and subjects were instructed to “say the name of the color, as fast and accurately as possible; do not read the word.” The primary manipulation concerns the *congruency* between the meaning of the text and the hue in which it is rendered. A colored word is either “congruent”, such that the hue corresponds to the hue the text expresses (e.g., “RED” in red font), or “incongruent” (e.g., “GREEN” in blue font).

The Stroop dataset within the DMCC project is highly amenable to assessing test-retest reliability due to the extensive amount of repeated measures acquired. Each subject completed several hundred trials of this task within each of six scanning sessions, administered on different days. By design, these six scanning days were clustered into two “repetitions” of three sessions – an initial “test”, then subsequent “retest” repetition. Across subjects, the time between repetitions spanned 25–1567 days (median = 209, IQR = 334), while the time between sessions, within each repetition spanned 1–70 days (median = 16.5, IQR = 21.5). Assessing test–retest reliability across these repetitions thus reflects a relatively challenging benchmark for consistency.

These sessions were largely similar within each repetition. Each consisted of two scanning runs of approximately 12 minutes that contained a minimum of 108 trials, with inter-trial intervals sampled with uniform probability from one, two, or three TRs (1.2, 2.4, or 3.6 s). The same set of eight words, and set of eight corresponding hues, were used in each session as stimuli. Yet by design, the sessions also subtly differed in ways that influenced the size of the behavioral congruency effects that they elicit. Although these manipulations were conducted to investigate questions outside of the scope of the present study, we exploit them in our test–retest reliability analyses. In what we have referred to as the “baseline” type of session (216 trials), the first session within each repetition, behavioral congruency effects were maximized (Braver et al., 2021). This maximization was due to the low frequency of incongruent relative to congruent trials (33%; Logan and Zbrodoff, 1979). The other two “proactive” and “reactive” sessions (216, 240 trials), which were counterbalanced in order across participants, manipulated expectancies regarding incongruent trials. The expectancy manipulations were accomplished by increasing the relative percentage of incongruent versus congruent trials, either in a session-wide manner (to 67%, proactive session), or selectively for specific colors (while keeping the session-wide percentage at 40%, reactive session). The theoretical reasons for the manipulations are described in detail within Braver et al., 2021. We exploit the robust congruency effects elicited by the baseline session in our analyses, by using this session in particular to evaluate the test–retest reliability of our univariate fMRI measures (see Section 2.6).

### 2.4 Image acquisition and preprocessing

The fMRI data were acquired with a 3T Siemens Prisma (32 channel head-coil; CMRR multiband sequence, factor = 4; 2.4 mm isotropic voxel, with 1.2 s TR, no GRAPPA, ipat = 0), then subjected to standard fMRIPrep pipelines (Esteban et al., 2018, 2020). As part of the preprocessing pipeline, data were projected into surface space (fsLR8K) once, were then smoothed (with Gaussian kernel of FWHM = 4 mm), and were divisively normalized to reflect percent signal change relative to the timeseries mean. These pipelines were implemented in a Singularity container (Kurtzer et al., 2017) with additional custom scripts used to implement file management. More details on the preprocessing and pipeline are available at https://osf.io/6p3en/ and (Etzel et al., 2022). Container scripts are available at https://hub.docker.com/u/ccplabwustl.

### 2.5 Timeseries models

To generate single-trial estimates of the fMRI activation, we summarized the minimally preprocessed fMRI timeseries using a simple “selective averaging” model. Specifically, we averaged together the fourth, fifth, and sixth observations (i.e., time-points in TR units) after the onset of a Stroop stimulus (i.e., corresponding to a window of 4.68–7.08 s post-stimulus onset, after accounting for stimulus presentation delays and slice-time correction). Prior to selective averaging, we detrended the timeseries for a given participant and repetition with 5-th order polynomials per run (with order selected by 1 + floor(*D/*150), where *D* is the duration in seconds of a run), as well as with 6 motion parameters concatenated in time over both runs. This detrending was performed via 3dDeconvolve in *AFNI*. Additionally, trials that had a TR within the averaged window with frame-wise displacement > 0.9 mm were censored.

We opted to use this selective averaging procedure due to its simplicity and because the variance of the estimates are not diminished by collinearity within the design matrix (c.f., GLM-based approaches that simultaneously model all trials with individual regressors, such as a “least-squares–all” approach; Mumford et al., 2012). In this way, the goal of using selective averaging is similar to that of other highly popular methods, such as “least–squares separate” (Mumford et al., 2012), in that some degree of bias is tolerated with the hope of reducing variance. But, because the degree of bias can be difficult to identify or control within selective averaging (an issue it shares with least-squares–separate), it was critical for us to validate this method. We therefore conducted several analyses (reported in Supplemental Figures 9 and 10) to confirm whether this method can yield estimates interpretable as single-trial activation amplitudes, and whether these estimates are highly similar to those from other, widely used GLM-based modeling methods. Results confirmed these validation checks, suggesting that our results and inferences are likely robust to these concerns.

### 2.6 Spatial brain activity models

The selective averaging procedure described within Section 2.5 furnishes a set of activation estimates per trial and vertex. We used the Schaefer atlas to parcellate these estimates into a set of *N* data matrices **X** of size *T* × *V_n_* for brain region *n* ∈ 1*, …, N*.

After we centered these estimates (see section below), we used two different methods of summarizing the spatial patterns within each region: a univariate model and a multivariate model. Each of these models aggregates information that is spatially distributed across all vertices within a given region. Thus, we refer to them as “spatial models” (equivalently, they could be referred to as “spatial filters”). In particular, these spatial models perform linear dimensionality reduction, in which the spatial pattern on each trial is projected onto a single dimension, yielding a single summary score, per trial and region. In other words, both univariate and multivariate models *m* ∈ {univ, multiv} compute a weighted sum across vertices:

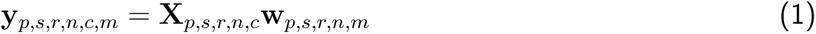

where **y** is a vector of length *T* that holds the projections for each trial.

The key difference between these models lies in the nature of the weights, **w**. In the univariate spatial model, the weights are all positive and equal across vertices:

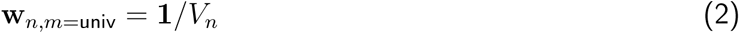

where **1** is a vector of length *V_n_* whose entries are all ones. This projection amounts to estimating the spatial mean.

In the multivariate spatial model, the weights are estimated such that they optimize a criterion. We used linear discriminant analysis (LDA; Fisher, 1936; Hastie et al., 2009, which finds the weights that maximize, within the projection scores, the variability *between the condition means* relative to the variability *within the conditions*. In other words, weights are learned that suppress trial-level variability relative to trial-averaged variability in the scores. Notably, this criterion aligns well with our goal of obtaining a favorable *trial/subject* variability ratio. Such weights are provided by the linear discriminant function within LDA:

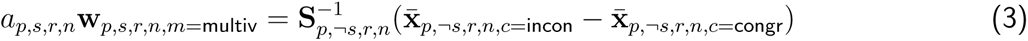

where 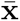 is the mean vector of activation estimates, averaged over trials and of length *V_n_*; **S** is the mean within-condition covariance matrix, averaged over conditions and of size *V_n_* × *V_n_*; and *a* is the Euclidean norm of the expression on the right-hand side, so that **w** is unit length.

The covariance matrix **S** is a critical part of this operation, in that it incorporates information regarding not only the amount of trial-level variability in each vertex, but also how this variability co-varies with each of the other vertices within the region. In this way, these vertex-wise scaling factors enable LDA to squash unreliable spatial dimensions and expand reliable ones.

The multivariate models were fitted in leave-one-session-out cross-validation, in which the weights for a given session *s* were estimated using concatenated data from the other two sessions, ¬*s*. To provide a moderate amount of robustness to the model, we regularized **S** by shrinking the off-diagonal values towards zero by a fixed amount of 25% (*cf.* Diedrichsen et al., 2016). We implemented LDA using the *rda()* function from the R package *klaR* (Weihs et al., 2005).

#### Centering

Prior to fitting the spatial models, the trial-level activation estimates from the timeseries model were centered. In particular, the values in each vertex were centered at their mean of condition means within each run. This centering is also known as “mean pattern” or “cocktail” centering (Misaki et al., 2010; Walther et al., 2016). The goal of implementing this centering was to reduce nuisance variance that resulted from global changes across different scanning runs and sessions (i.e., “global”, in the sense of changes that impact all conditions equally within each run). While this form of centering has been criticized for rendering linear correlations difficult to interpret (eGarrido et al., 2013), the concern does not apply in the present case with an LDA-based decoder (which implements a centering operation by default; see Misaki et al., 2010; Walther et al., 2016). To avoid imposing systematic differences between runs via centering, we estimated the mean using only the specific subset of trials that were present in each scanning run with matched stimulus features across runs and with balanced proportions of congruent and incongruent conditions within each run (see King et al., 2019 for a similar approach). Finally, we implemented these steps prior to fitting both multivariate and univariate spatial models, to ensure that the resulting model outputs **y**, differ only due to differing weights **w**, and not differing input data **X**.

#### Stratified random undersampling

To minimize bias in decoding results caused by class imbalance and other confounding factors, we used a stratified random undersampling algorithm to generate the training sets for LDA. By randomly sampling subsets of trials within our training set without replacement, we built a collection of 100 smaller training sets that contained an approximately equal number of trials *k* of each word and color within both congruent and incongruent conditions of each scanning run. We selected *k* as the minimum number of occurrences in any given combination of these factors within a given scanning run. Due to slight differences in trial balancing, the exact number *k* differed across across runs and sessions: *k* ∈ {4, 5} in the baseline session, *k* ∈ {1, 2} in the proactive session, and *k* = 6 in the reactive session. Note that these differences should not lead to systematic biases in the results, as we pool over both runs of two sessions when training decoders, so that the imbalance is systematically washed out.

### 2.7 Reliability models

After the fMRI activity patterns within each parcel were summarized into univariate and multivariate projection scores per trial, **y**, the test-retest reliability of these projections was estimated. We compared two different methods of estimating test–retest reliability: the Intraclass Correlation, which is a standard “summary statistic” method, and correlations derived from hierarchical Bayesian models. We refer to both of these methods generally as “reliability models”.

We fitted both summary statistic and hierarchical Bayesian reliability models similarly and independently for each of the 400 Schaefer-atlas brain regions, and each type of spatial model (univariate, multivariate). This enables us to compare the independent and interactive effects of spatial and reliability modeling methods, similar to decomposing an effect into main effects and interactions.

We scope our test–retest reliability analyses, however, exclusively on projections from the baseline session. This decision was made to streamline reporting of our results. There is strong reason to expect that individual differences in the Stroop effect were maximized within this session (see Section 2.3; Braver et al., 2021). Consequently, this decision likely works *against* our key hypothesis that multivariate models improve properties of test–retest reliability relative to univariate models. This is because the test–retest reliability of multivariate projections is not only dependent on baseline-session data, but also data from the other two sessions (proactive and reactive), as those sessions constituted the training set (in cross-session cross-validation). By contrast, univariate projections within the baseline session do not have this dependency. In other words, focusing on baseline-session data not only allows us to simplify our results, but it gives univariate models their “best-shot” in comparison against multivariate models. Thus, any improvements we see in properties of test–retest reliability due to multivariate models provides strong evidence for their superiority.

#### 2.7.1 Summary-statistic models

The Intraclass Correlation Coefficient (ICC) is a standard way to measure reliability. In particular, a widely used form quantifies the consistency instead of absolute agreement between two repeated measures within the same group of participants [ICC(3, 1) in Shrout and Fleiss, 1979]. We used this measure as a “summary-statistic method” of estimating reliability.

First, we averaged the trial-level activation estimates over trials, forming one summary score per subject, repetition, and condition (incongruent, congruent): 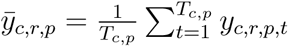. The contrast of interest is the Stroop effect, that is, the difference between means of incongruent versus congruent trials, which we denote as 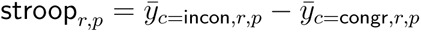.

Then, at the population level, we modeled individual variability in the Stroop effects with a set of Gaussian distributions:

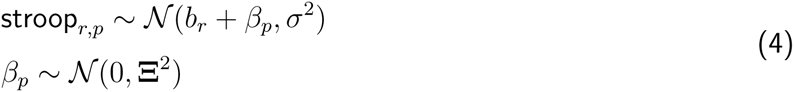

where *b_r_* is the “fixed” (population-level) Stroop effect associated with repetition *r*, and *β_p_* is the “random” (individual) Stroop effect associated with participant *p*, on average across sessions. The covariance in participants’ Stroop effects across test and retest repetitions is the single off-diagonal entry of the covariance matrix **Ξ**^2^, which we denote as *ξ*^2^ for simplicity. Under this formulation 4, the intra-class correlation coefficient is defined as the proportion of variance:

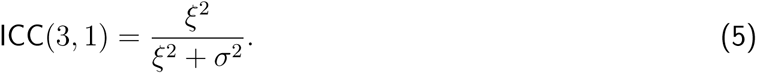

Alternatively, it can be shown that ICC(3, 1) is the same as the Pearson correlation coefficient between the Stroop effects of both repetitions over the participants *p*:

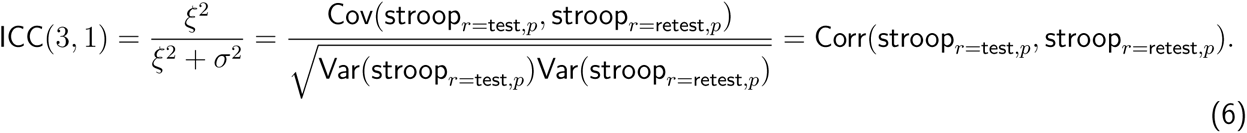

#### 2.7.2 Hierarchical Bayesian models

In the hierarchical Bayesian method of estimating reliability, projections are modeled at the trial level, and the amount of variance at different levels — such as across trials versus across subjects — is separately captured. Unlike intra-class correlation, a hierarchical Bayesian decomposition enables individual-difference correlations to be estimated in a manner that accounts for the impact of variance at each level.

For detailed formulations of our hierarchical models, see subsequent section on Hierarchical Model Structures. Here, we provide a simplified description. The structure of our hierarchical models generally followed prior work (Chen et al., 2021). For readers familiar with Wilkinson and Rogers, 1973 notation (e.g., *lme4* syntax). a key part of our model can roughly be expressed as

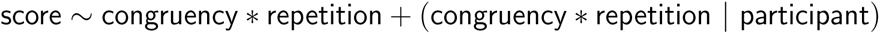

in which the non-parenthetical terms on the right-hand side denote the population (“fixed”) effects, while parenthetical terms denote individual-level (“random”) effects, and “score” on the left-hand side denotes the trial-level projection scores, **y**. The quantity of test–retest reliability is contained within the parameters associated with the individual-level effects.

Models were fitted using the R package *brms* with noninformative or weakly-informative hyperpriors automatically selected by *brms* (Bürkner, 2017). Parameters were estimated through Markov Chain Monte Carlo (MCMC) with four chains with 2000 iterations each (including 1000 warm-up iterations). The training time for each model (i.e., for univariate or multivariate contrasts of a single parcel in the baseline session) was approximately an hour, running on four threads on an Intel Xeon E5-2670 CPU (2.60 GHz).

To ensure that we used models well-fitted to our dataset, we assessed four models of varying complexity and compared them in estimates of their ability to predict new data points (see Supplementary Results, Model Comparison). These models varied in terms of the complexity of the covariance structures in the location and scale parameters of the *t* distributions (see section on Hierarchical Model Structures).

Due to computational constraints, this comparison was conducted in a subset of 32 parcels within the 400-parcel Schaefer atlas. These parcel locations were based on prior work that demonstrated a core set of regions associated with demands for cognitive control (Assem et al., 2020, within the Glasser atlas), and were independently selected in prior work (see Braver et al., 2021 Supplemental Materials https://osf.io/pa9hj for identifying corresponding regions within the Schaefer atlas). Expected log-predictive density with Bayesian leave-one-out cross-validation (ELPD LOO) was computed by *brms* through the *loo* package (Vehtari et al., 2024). A model of intermediate complexity was supported by this criteria in all 32 parcels for both univariate and multivariate approaches (Reduced Model 2; Equation 9). Another model fit statistic, widely applicable information criterion (WAIC), also supported this model (not reported). Therefore, we then separately fitted this model to data from all 400 parcels, selecting it as the basis of analyses reported in the Results.

##### Hierarchical Model Structures

###### Full model

The full or “maximal” model structure is denoted in Equation 7 and is explained in detail in the rest of this section.

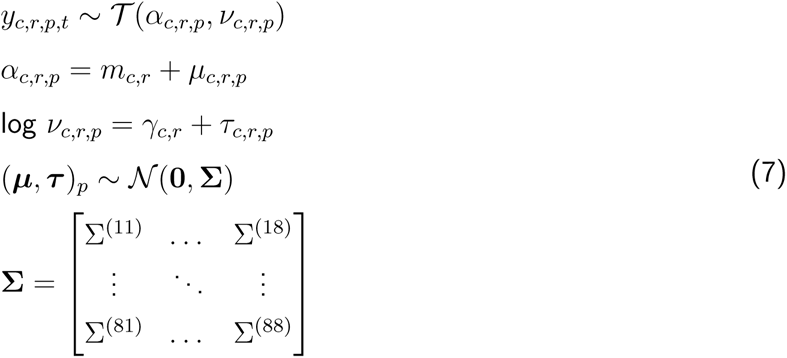

We begin by assuming the observed output of the spatial models, *y* (*cf*, **y** in Equation 1), is generated from a set of Student’s t-distributions, each of which are fully described by a location parameter *α* and scale parameter *ν*. Conceptually, these location and scale parameters of the *t*-distribution are analogous to the mean and standard deviation parameters of the Gaussian distribution. The *t*-distribution is used here instead of Gaussian, however, as its fatter tails provide greater robustness to outlying values (e.g., Chen et al., 2021). Importantly, the subscripts on *α* and *ν* indicate that individual parameters are estimated for each participant *p*, condition *c* ∈ {incon, congr}, and repetition *r* ∈ {test, retest} combination.

Note that the the condition and repetition factors are parameterized with a “flat” coding scheme (also sometimes referred to as a “no-intercept” dummy coding scheme). Under this scheme, a given participant’s *α* parameter coefficients specify their mean projections for each condition and repetition combination (e.g., incon within the test repetition).

Next, we assume that these location parameters *α_c,r,p_* are a sum of two components: *m_c,r_*, a population-level fixed effect, which all participants share; and *µ_c,r,p_*, a participant-level random effect, which indicates the deviation of the *p*-th participant’s score from the group score *m_c,r_*, within repetition *r* and condition *c*.

Similar to the location parameter, we decompose the (log of the) scale parameter into population and participant-level effects. We denote these effects with *γ_c,r_* and *τ_c,r,p_*, which are analogous to their location-parameter counterparts. In effect, these parameters enable the model to account for differing amounts of residual variability for each participant, repetition, and condition combination.

Finally, we assume that each participant’s eight location and scale parameters, now written as a single 8-element vector (***µ***, ***τ***), were sampled from a single multivariate Gaussian distribution with zero mean and covariance matrix **Σ**^2^. The covariance matrix **Σ**^2^ describes the linear relationships among these parameters over subjects.

Contained within **Σ**^2^ is the pivotal quantity of test-retest reliability: the correlation between test and retest repetitions in the Stroop contrast, denoted here as *ρ*. Due to the flat parameterization of the condition and repetition factors, however, *ρ* is not explicitly expressed as a single term in **Σ**^2^, but instead implicitly, as a linear combination of its rows and columns. To obtain *ρ*, we transform **Σ**^2^ by a 2-by-8 contrast matrix **W** that encodes this linear combination. Specifically, row *r* of **W** corresponds to the test or retest repetition, while column *j* corresponds to the *j*-th element of the participant-level random effect vector (***µ***, ***τ***). Where *j* corresponds to *µ*^(^*^c^*^=incon^*^,r^*^)^, *W* ^(^*^rj^*^)^ = 1; where *j* corresponds to *µ*^(^*^c^*^=congr^*^,r^*^)^, *W* ^(^*^rj^*^)^ = −1; elsewhere, *W* ^(^*^rj^*^)^ = 0. Applying this contrast to the random-effect covariance matrix, **WΣ**^2^**W**^⊺^, and dividing the resulting covariance element by the product of the standard deviations, yields the test-retest correlation in the Stroop effect, *ρ*.

###### Reduced model 1: independent location and scale (ILS)

The first simplification to the “full model” that we considered was omitting the random-effect covariances between the location ***µ*** and scale ***τ*** parameters. This simplification led to a model with the same general form as in Equation 7, except location ***µ*** and scale ***τ*** parameters were assumed to be generated by independent distributions, with independent covariance matrices 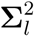, for *l* ∈ {locat, scale}.

From Equation 7, the reduced terms are as follows:

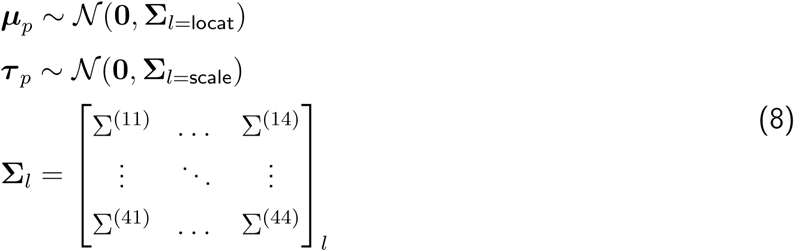

###### Reduced model 2: independent location and scale, symmetric covariance structure (ILS Sym)

We further simplified the model with independent location and scale (Equation 8) by additionally assuming that the covariance structure was symmetric between repetitions (Chen et al., 2021). Under a symmetry assumption, the covariance between different conditions within different repetitions is constrained to be equal across permutations of repetitions. For example, a symmetric structure would entail that Cov(incon test, congr retest) equals Cov(congr test, incon retest). To parameterize this covariance structure, we use a different coding scheme to parameterize the condition factor than used in Equation 7. Now, we use a contrast coding scheme, *c*′ ∈ {mean, stroop}, such that, for a given participant *p* and repetition *r*, *α_c_′*_=mean_*_,r,p_* represents the mean of incongruent and congruent condition means, and *α_c_′*_=stroop_*_,r,p_* represents the difference between incongruent and congruent means (i.e., the mean Stroop contrast). Under the symmetry assumption, the mean of incongruent and congruent conditions is independent from the mean Stroop contrast over subjects (Chen et al., 2021).

From Equation 7, the reduced terms are as follows:

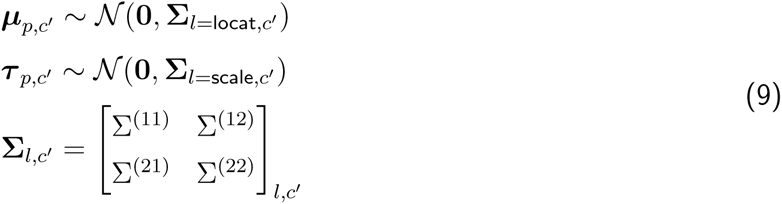

Note that ***µ*** is now a vector with 2 elements, corresponding to each repetition *r*. Under this contrast coding scheme, it is now no longer necessary to apply a contrast matrix to **Σ** to obtain the test-retest correlation in Stroop effects *ρ*.

###### Reduced model 3: independent location and scale, symmetric covariance structure, homogeneous scale (Homog.)

Finally, the simplest model we considered was a reduction of the model with symmetric covariance structure (Equation 9), in which we additionally assumed that the scale of the residuals, *ν*, was constant across all conditions, repetitions, and participants.

From Equation 7, the reduced terms are as follows:

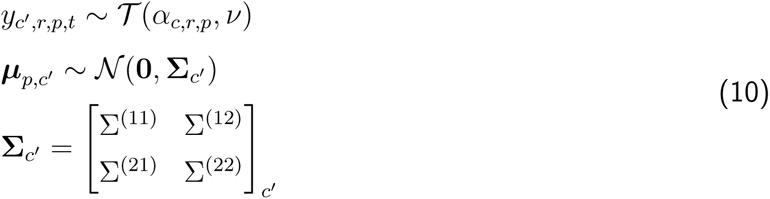

Note that, although this homogeneous-variance assumption is likely unrealistic, it is typically made by default: for example, it is required by popular hierarchical-modeling libraries such as *lme4*.

##### Analysis of Hierarchical Model Parameters

###### Population-level Stroop effect

The posterior distribution (MCMC samples) of the “fixed” or population-level effect of congruency for each repetition was extracted from the models through the *fixef()* function of *brms*. These estimates correspond to the *m_c_′*_=stroop_*_,r_*in Equation 9. Samples from this distribution were averaged over repetitions *r*. Then, a *t*^+^ statistic was computed as the ratio between the mean and standard deviation of the posterior distribution.

###### Point estimates and precision of test–retest reliability

The posterior distribution of TRR was extracted through the *VarCorr()* function of *brms*.

In the main text, we primarily use the *maximum a posteriori* (MAP) estimate to represent the central tendency of the posterior TRR distribution. We selected this measure to be consistent with closely related prior work (Chen et al., 2021) and for its convenient properties. Namely, the MAP captures the most likely value of the posterior, which is particularly useful if the distribution has a prominent peak representing high probability density. The MAP is also informative when the distribution is skewed, as often occurs when test-retest reliability values approach boundary limits.

We also report results with posterior means within the Supplemental Materials. To compute the posterior mean, TRR values were *artanh* transformed, averaged over samples, then *tanh* transformed. Posterior medians were very highly correlated with posterior means, so we do not report them. In general, our results and inferences do not depend strongly on the choice of summary measure.

The dispersion of the posterior distribution was summarized by the precision, which is the inverse of the standard deviation.

###### Variability ratio

We quantified the relative magnitude of trial and individual-level variability in the Stroop effect contrast as a ratio of standard deviations. As there are multiple standard-deviation estimates at each level (i.e., one per repetition), we aggregated them before computing the ratio, and we did this via the geometric mean. From Reduced Model 2 (Equation 9), the particular standard deviations are computed as follows: the trial-level (within individual) standard deviation is given by the population-level mean of the scale parameter *γ_c_′*_=mean_, averaged over the two repetitions,

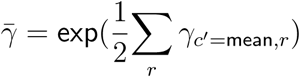

whereas the individual-level (between individual) variability is given by the standard deviation of Stroop effects, averaged over the two repetitions,

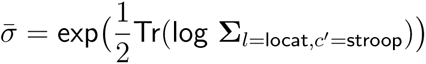

where Tr is the matrix trace. (Note that *γ* is estimated by the model on the log scale, whereas **Σ** is not.) The variability ratio was computed as log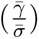 separately for each MCMC sample, then summarized across samples by the mode of the distribution.

## 3 Results

### 3.1 Pronounced univariate activation in fronto-parietal networks at the population level

To provide a basis for assessing reliability of cognitive control-related BOLD responses at the level of individuals, we first characterized BOLD responses that were co-localized across individuals within particular brain regions. Namely, we assessed which cortical regions (parcels within the Schaefer 400 17-Network atlas) increase BOLD activity on more demanding incongruent Stroop trials, relative to less demanding congruent trials, generally across subjects, as such changes in activity are expected to be exhibited by a region involved in cognitive control. This contrast was implemented by first spatially averaging the BOLD signals across vertices within each parcel separately on each trial, then estimating the population-level change in activation within a hierarchical Bayesian model (Reduced Model 2 in Method, Equation 9). Hereafter, we refer to this contrast as a “univariate activation” contrast, as it relies on a spatially univariate (i.e., uniform) averaging of signals from a given region. Figure 2 displays the results of this analysis.

**Figure 2:**
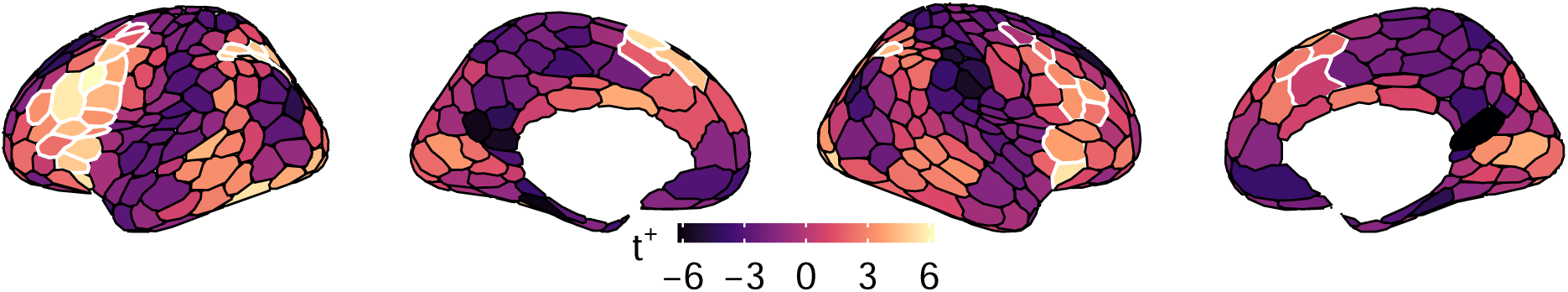
Population-level univariate activation to Stroop task demands. Per brain parcel, *t*^+^ statistics show the sign and magnitude of univariate activation. These statistics were computed by estimating the spatially averaged difference of BOLD activity estimates on incongruent versus congruent trials, then dividing this estimate by its standard error, estimated through a hierarchical Bayesian model. Regions of interest, which are outlined in white borders, were defined using a larger portion of the Dual Mechanisms of Cognitive Control dataset.

To illustrate the continuous magnitude and sign of the change in activation across parcels, we depict the effect size with a statistic we refer to as *t*^+^, which incorporates both the mean level and amount of uncertainty of activation change (note, however, that although similar in form, the definition of *t*^+^ does not straightforwardly translate to the frequentist’s t-statistic). Using this contrast, prominent increases in activation were observed in frontal and parietal cortex, particularly within the left hemisphere (Figure 2), in a pattern that is strongly consistent with extensive prior findings (e.g., Assem et al., 2020; MacDonald et al., 2000).

To complement this continuous measure, we also employed a dichotomous measure, by using “regions of interest” (ROIs) based on the larger Dual Mechanisms of Cognitive Control dataset from which this subset of data was taken. Specifically, in Braver et al., 2021 a set of 35 parcels were identified from a larger sample of participants (N = 80) that showed consistent activation according to cognitive control demands, and which included the Stroop effect contrast, but also parallel contrasts across three additional tasks studied within that report. Compared to using the present sample and task alone, defining ROIs on the basis of the larger Dual Mechanisms dataset likely yields more accurate definitions of core brain regions associated with cognitive control across multiple tasks. These ROIs included brain areas classically associated with cognitive control, such as mid-lateral prefrontal and posterior parietal cortices (Figure 2, white borders), and which most prominently belonged to the fronto-parietal control network.

In subsequent analyses, we will use both of these complementary statistics (*t*^+^, ROI definitions) to illustrate relations between univariate activation at the population level, and properties of other neural measures at the individual level, such as test–retest reliability.

### 3.2 Hierarchical modeling reveals highly reliable estimates at the individual level

Having established that univariate contrasts reveal highly robust activity changes on average *across* individuals, we next asked whether these same univariate contrasts are also highly consistent *within* each individual, across repeated testing sessions, which in our sample were separated by several months to years of intervening time. In other words, what is the test–retest reliability of fMRI activation to Stroop-task demands?

To provide a comprehensive assessment of test–retest reliability, we used two different approaches for estimating reliability and compared their results. The first approach was via a widely used method, the intra-class correlation, which relies on a two-stage procedure in which “summary statistics” are computed, then correlations in these statistics are subsequently estimated. The second was via a hierarchical Bayesian modeling approach. Using the same models we fitted to estimate the population-level effects separately in each brain parcel (i.e., in Figure 2), we also estimated the posterior distribution of test–retest correlations within individuals. Following prior work Chen et al., 2021, to summarize these posterior correlations with a single value, we used the maximum *a posteriori* probability (MAP) estimate (i.e., the single correlation value at which the density of the posterior probability is maximal).

Figure 3, top, depicts the results of both of these approaches, as applied to individuals’ univariate activation contrasts. When estimated through a summary statistic approach, test–retest reliability is quite modest, reaching maximum values of only *r* ∼ 0.5, primarily within our ROIs (white borders), which tended to contain strong population-level effects. The results from the hierarchical Bayesian model were quite different. As revealed by these models, test–retest reliability was often close to maximum, again prominently within ROIs, but also within a much broader set of regions. In fact, the number of regions with conventionally “high” reliability (*r >* 0.7) dramatically increased from 3 to 186, indicating that nearly 50% of the brain parcels were highly reliable.

**Figure 3:**
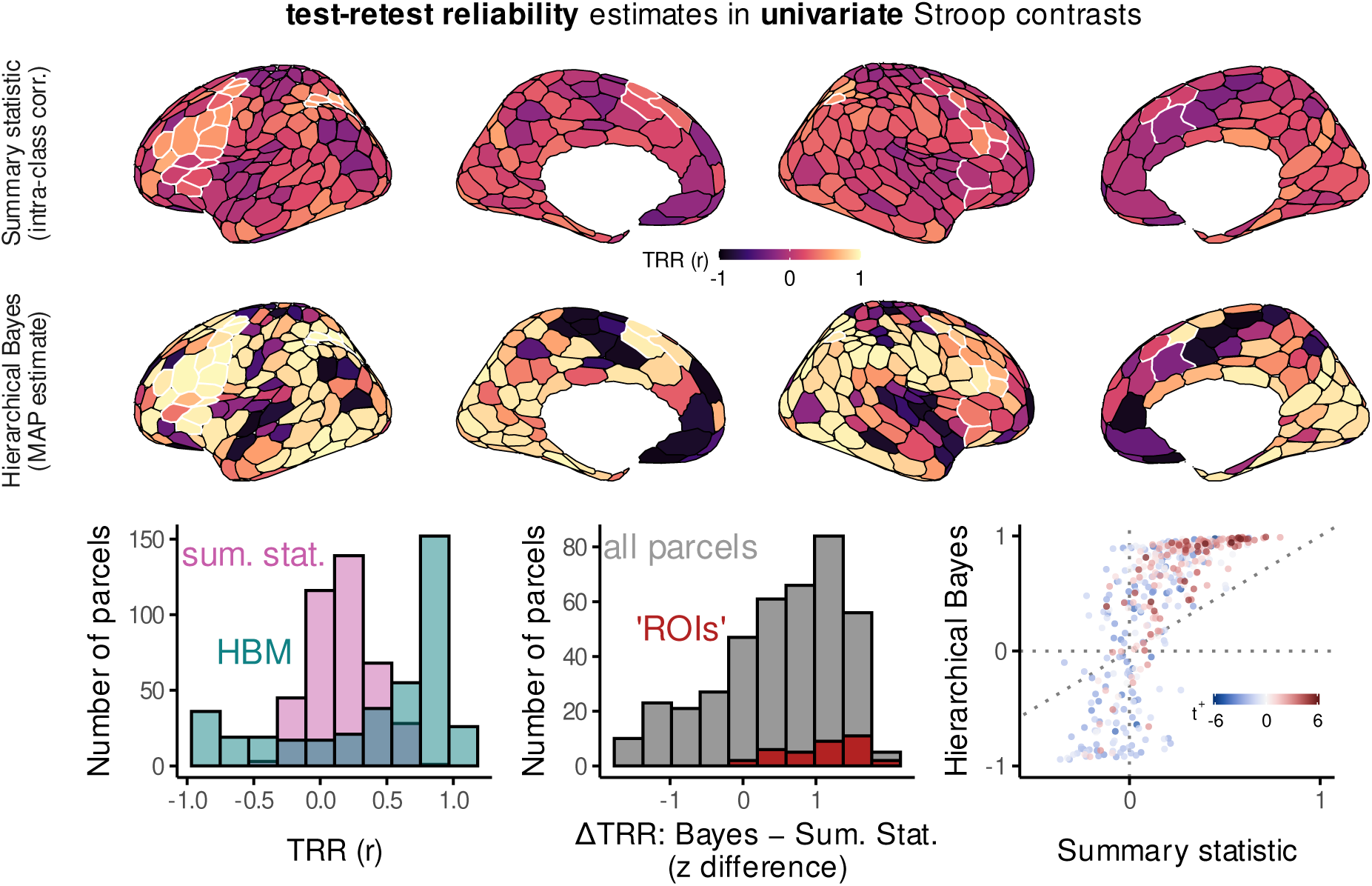
Test–retest reliability estimates in univariate Stroop contrasts. **Surface plots at top**, Test–retest reliability (TRR) correlation coefficients (r) estimated through “summary statistic” (upper row) or hierarchical Bayesian models (bottom row). To summarize the posterior correlation distributions in the hierarchical models, we used the maximum *a posteriori* (MAP) estimate. White borders illustrate regions of interest (ROIs) identified in a prior report (Braver et al., 2021). **Bottom left**, Histogram of the distribution of these test–retest correlations across all cortical parcels (hierarchical Bayesian models or HBM, pink; summary statistic, sum. stat.). **Bottom middle**, Histogram of the difference in test– retest correlations between hierarchical Bayes MAP estimates minus summary-statistic estimates, over all cortical parcels and also in ROIs. Prior to subtraction, correlation coefficients were z-transformed (i.e., inverse hyperbolic tangent). **Bottom right**, Scatterplot of the relation between test–retest correlation estimates from summary statistic models (x axis) and hierarchical Bayesian models (y axis), over all cortical parcels (points). The color scale illustrates the sign and magnitude of the population-level univariate Stroop effect contrast (*t*^+^, defined in Figure 2). Dotted lines illustrate x and y intercepts, as well as the unity line. **Supplemental Materials**, For an analogous figure that displays posterior mean TRR instead of MAP, see Supplemental Materials Figure 18.

By directly contrasting the summary-statistic and hierarchical correlations within each parcel, the differences between the estimates can be explicitly illustrated. Two findings here are notable. First, the change in reliability was most prominent for regions in which the intra-class correlations was already distant from zero. For regions with positive intra-class correlations, hierarchical modeling dramatically improved reliability, while for regions with negative intra-class correlations, hierarchical modeling reduced reliability. For regions with near-zero intra-class correlations, little change was revealed. This finding is consistent with prior theoretical and empirical demonstrations that, compared to hierarchical Bayesian models, intra-class correlation multiplicatively underestimates test–retest reliability (Chen et al., 2021). Second, improvements in reliability revealed by hierarchical modeling also tended to be particularly large in regions with strong population-level effects, that is, fronto-parietal regions which were consistently activated to Stroop-task demands across subjects. Consequently, the same experimental contrasts used to identify regions that are involved in cognitive control, generally within the population, can also be used to individuate people, in terms of the extent to which their neural responses within these regions were modulated by the task (e.g., Braver et al., 2010).

We then sought to characterize reliability of an alternative contrast, of multivariate activation. Similar to the univariate activation contrast, the multivariate contrast we used implemented a linear transform of spatial patterns of brain activity, in which activity evoked by easier congruent trials was subtracted from activity evoked by more difficult incongruent trials. But instead of a uniform weighting of signals within a region of interest, the multivariate contrast relied on an optimal estimation of the weighting vector (see Method). Consistent with prior results, these multivariate contrasts were highly successful in discriminating incongruent from congruent trials’ activation patterns within many brain regions across the cortex, on average across individuals (Supplemental Figure 17; Braver et al., 2021; Woolgar et al., 2016). But how consistent are these multivariate contrasts within individual, across repeated testing sessions? The reliability of such a contrast has not yet been established.

In general, findings were quite similar to univariate contrasts: reliability was dramatically improved by hierarchical modeling, and the most reliable estimates were revealed in many of the same regions (Figure 4). In several regions, however, the gains revealed by hierarchical modeling were not quite as extreme as observed univariate contrasts. We will return to this observation in the Discussion.

**Figure 4:**
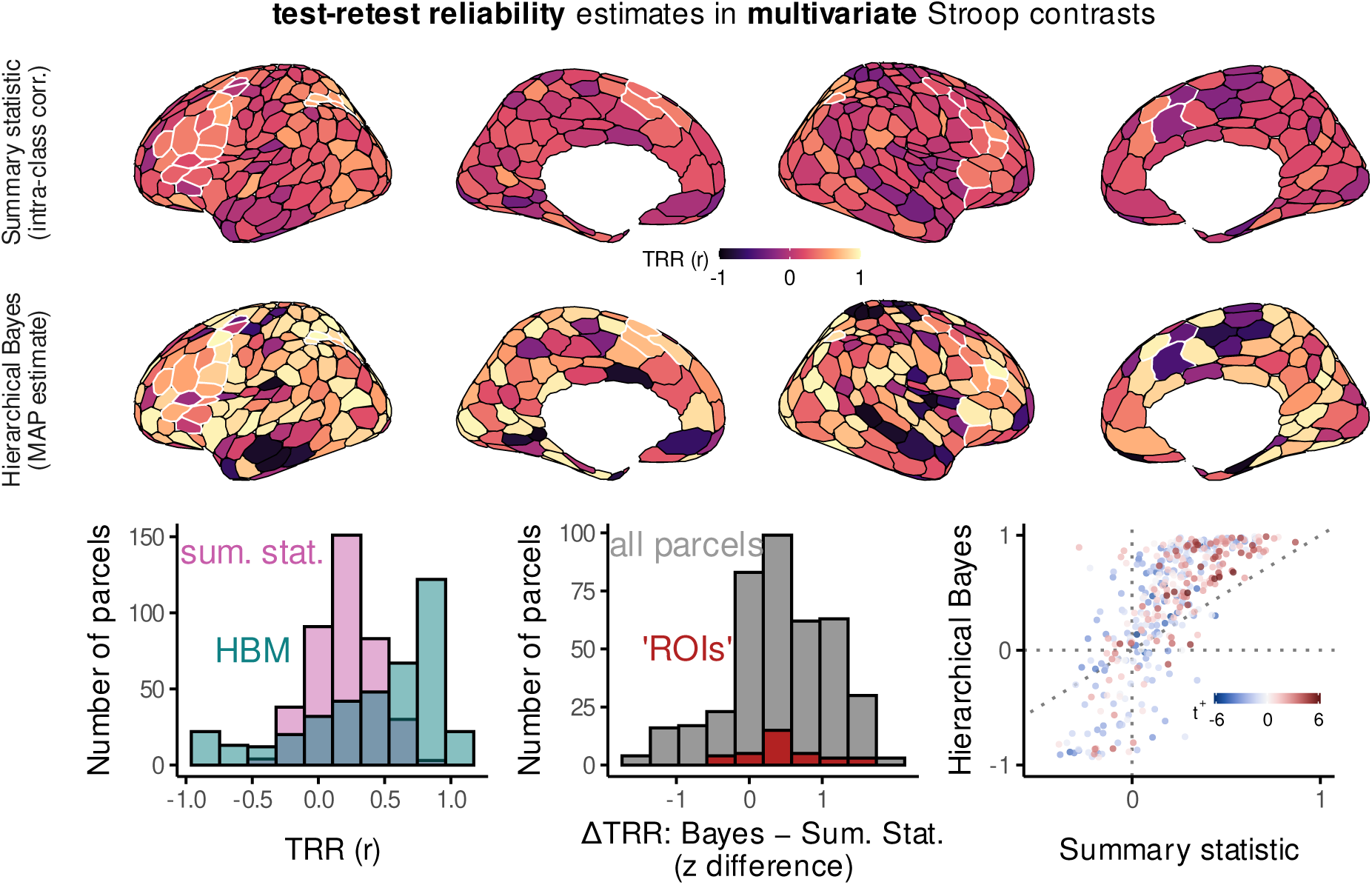
Test–retest reliability estimates in multivariate Stroop contrasts. All plots are analogous to those in Figure 3. For an analogous figure that displays posterior mean TRR instead of MAP, see Supplemental Materials Figure 18.

Collectively, these findings demonstrate that, both univariate and multivariate fMRI contrasts can be highly consistent within individual, at least when basing the estimation on the most likely (MAP) estimates from hierarchical models.

### 3.3 Imprecision limits interpretation of individuals’ univariate activation estimates

The results summarized in Figure 3 reflect the single most likely test–retest correlation value per brain region, as provided by the MAP estimator. While a convenient summary, this estimator conveys no information about *how much more likely* the MAP estimates are, relative to other outcomes in which reliability is lower. For instance, if the model assigns almost as much probability to a pattern of results in which reliability is considerably lower than in Figure 3, the prior results should not be well trusted. Evaluating these possibilities requires assessing the *precision* or *uncertainty* in the test–retest reliability estimates. Such an assessment is naturally supported within a Bayesian modeling framework, through analysis of the tails of the posterior distribution of test–retest correlations.

We found that, even as the single most likely test–retest reliability estimates of many regions approached a maximal value of one, in these same regions, the hierarchical model also assigned relatively high probabilities to correlation values that were quite low, and even below zero. We depict this uncertainty in Figure 5 (upper, left), in which we plot the MAP posterior test–retest correlation estimates against their lower-bound confidence intervals (5th percentile of the posterior, 5%-ile). From this view, only a handful of parcels now have a combination of both high reliability (> 0.7), and high certainty in the univariate reliability estimate (> 0). The sparsity of jointly reliable and certain univariate measures can also be seen in Figure 5 (lower, left) when thresholding the MAP correlation estimates based on their lower bounds. These findings converge with prior work (Chen et al., 2021).

**Figure 5:**
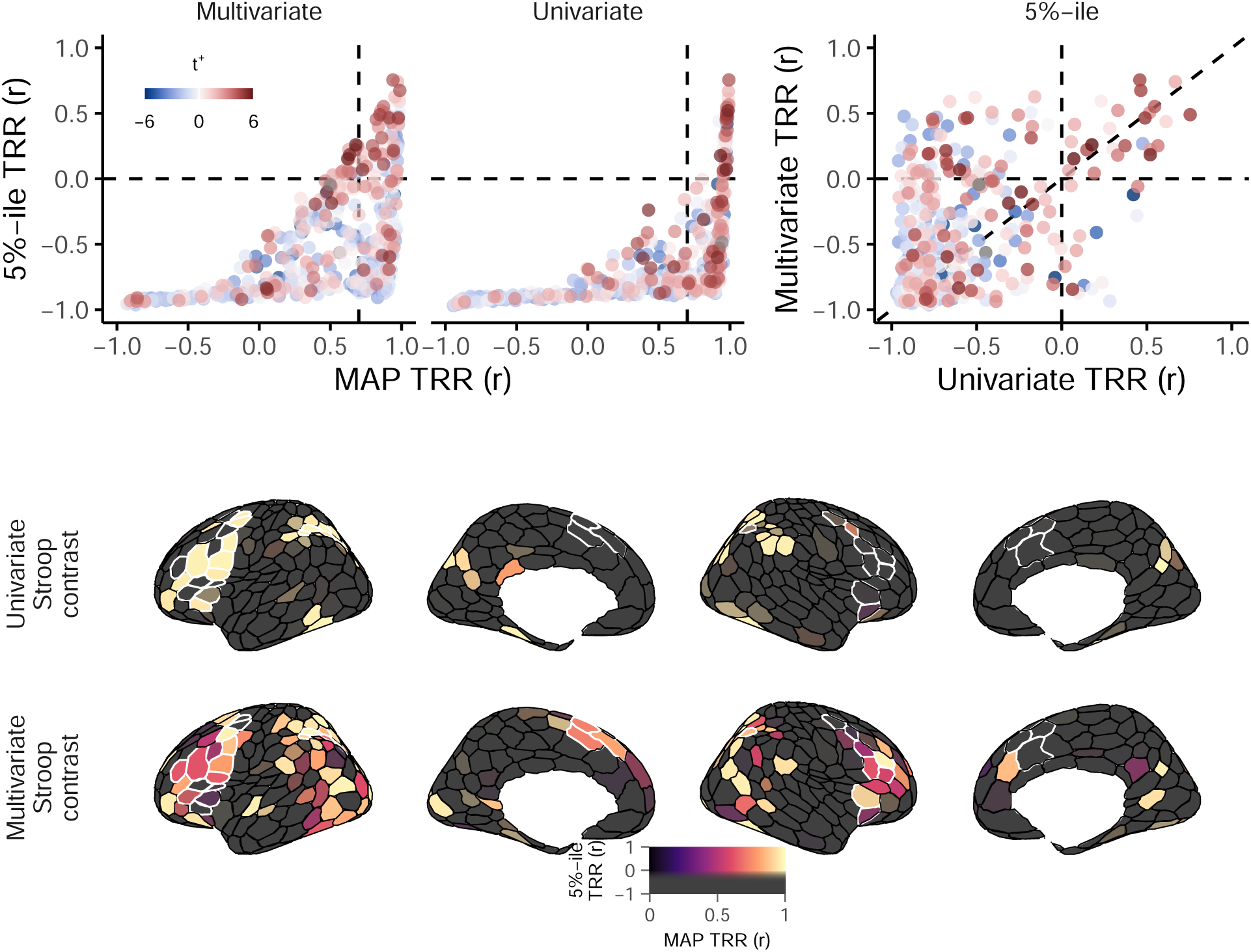
Lower-bound estimates of test–retest reliability correlations. **Upper left**, For each brain parcel (points), the MAP posterior test–retest correlation estimates (x axis) are plotted against their lower-bound estimate, the 5th percentile of the posterior (y axis). Although many parcels have near-maximal univariate reliability estimates, much fewer have confidence intervals that do not include zero. **Upper right**, The lower-bound on multivariate reliability (y axis) tends to be higher than that of univariate reliability (x axis; more points above diagonal, particularly when negative on x axis). **Upper**, All points are colored by their population-level univariate activation statistic (*t*^+^, see Method). **Lower**, Test–retest correlations for each cortical region, thresholded by their lower bounds. The black-to-hot dimension in the colormap indicates the MAP posterior test–retest correlation for univariate (top) and multivariate (bottom) contrasts. The saturated-to-unsaturated dimension in the color-map reflects the lower bound of the test–retest correlation estimate (95^th^ percentile). A soft threshold is applied to the color-map, so that only parcels that were estimated to have positive reliability with high certainty are displayed with full saturation (more colorful or deeper black), while those with increasingly negative lower bounds are displayed with quadratically decreasing saturation (more grey; Taylor et al., 2023). For an analogous figure that displays posterior mean TRR instead of MAP, see Supplemental Materials Figure 19.

In analyses focused on individual difference questions, observing such a high degree of uncertainty in individual-level variables would severely complicate inferences, weakening conclusions drawn about any relations observed.

### 3.4 A hierarchical-multivariate approach yields highly reliable and precise measures of individuals

Are multivariate contrasts subject to similar amount of imprecision as univariate contrasts? Given that multivariate contrasts optimize a different criterion than univariate contrasts (see Method), we suspected they may yield more precise individual-level measures. A hint to the answer is already provided in Figure 5. Although the central tendencies tend to be somewhat numerically lower, compared to univariate contrasts, multivariate contrasts identify over twice as many regions (80 versus 39) with high certainty (> 95% probability) of positive reliability. Of note, this category also included over half (19*/*35) of the ROIs. For instance, several regions in lateral PFC (particularly right hemisphere), posterior parietal cortex, and superior frontal cortex are identified as highly certain to have positive reliability, but only when the multivariate contrast is used. In Table 2, we list the 40 parcels with the highest (most positive) lower-bound estimates of test–retest reliability. Most of these parcels (26/40) are located within two fronto-parietal networks, Dorsal Attention A and Control A.

To provide a more comprehensive comparison, however, we calculated the precision of test–retest reliability separately in univariate and multivariate contrasts, then contrasted their magnitudes within each brain parcel. We used precision, that is, the reciprocal of the standard deviation of the test–retest correlation posterior, so that higher values indicate more precise estimates. (To ensure a sensitive comparison across a large range of values, we logarithmically transformed SDs prior to taking the reciprocal.) As illustrated in Figure 6, multivariate contrasts generally led to more precise individual-level contrast estimates.

**Figure 6:**
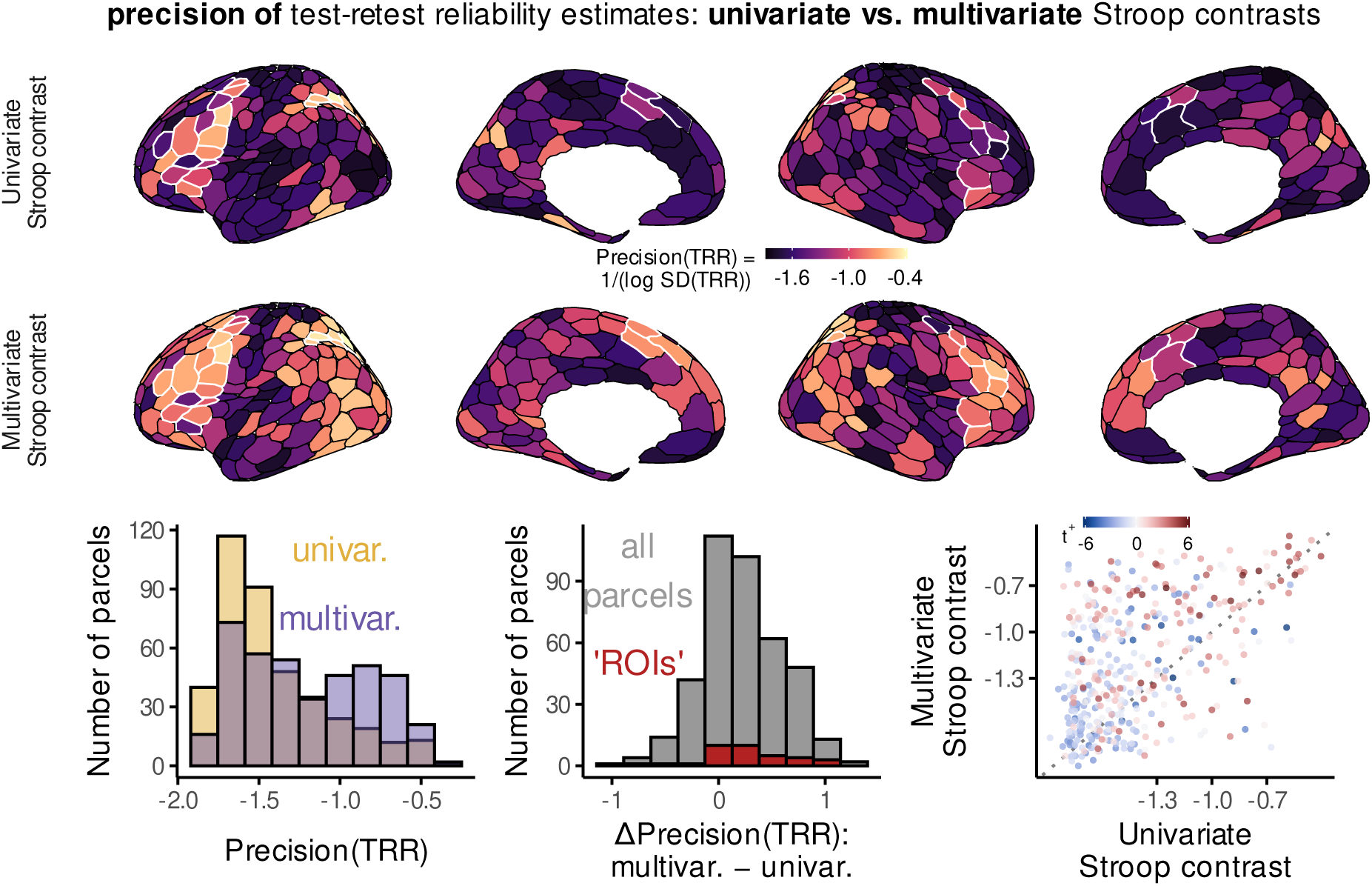
Precision of test–retest reliability estimates in univariate versus multivariate contrasts. We defined the precision of test–retest reliability correlations as the reciprocal of the log-transformed standard deviation across samples from the posterior distribution. Higher values indicate less variable (more certain) estimates of test–retest correlations. Aside from the difference in the statistic of interest, all plots are analogous to those in Figure 3.

We can now consider the joint impact of hierarchical and multivariate approaches on reliability of individual-level variables. For both univariate and multivariate contrasts, hierarchical modeling revealed that the central tendency of the posterior test–retest reliability distribution are often much greater than expected based on less-accurate summary statistic estimates (shown for multivariate contrasts in Figure 7 left panel, y axis). Yet relative to univariate contrasts, multivariate contrasts more strongly pulled the tails of the posterior distributions closer to their central tendencies (Figure 7 left panel, x axis). This not only increased the certainty in the plausible range of the parameter estimates, but also made these model results easier to summarize and interpret with point estimates. Such changes were relatively widespread, in that over 50% of parcels (233/400) show both improved reliability compared to summary statistic modeling (Figure 7), and improved certainty in individual-level estimates relative to univariate (Figure 7, left panel, upper right quadrant). Thus, this combination of methods can lead to a complement of benefits that has eluded prior work in this area: both *highly reliable* individual-level estimates, as well as *highly certain identification* of individual-difference associations.

**Figure 7:**
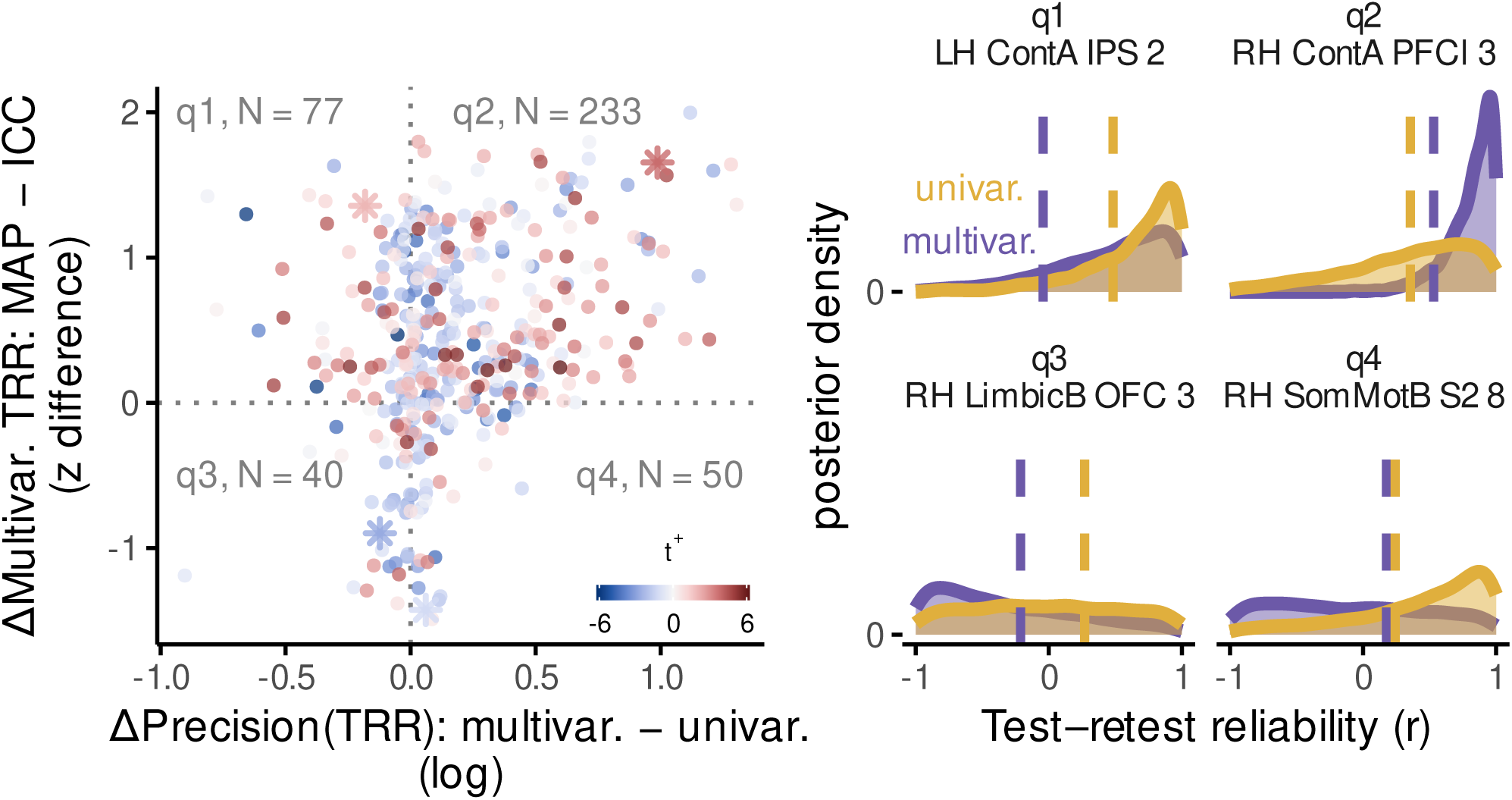
Joint impact of multivariate contrasts and hierarchical Bayesian modeling on individual-level correlation estimates. **Left**, Scatterplot of the joint benefits of multivariate and hierarchical modeling. The x axis depicts the increase in test–retest correlation precision associated with using a multivariate (multivar.) versus univariate (univar.) contrast (x axis). The y axis depicts results from multivariate contrasts only: namely, the increase in test–retest correlation strength revealed by hierarchical (maximum *a posteriori* estimate, or MAP) versus summary-statistic models (intra-class correlation coefficients, or ICC). “N=(#)” indicates the number of parcels in each quadrant. The color-map illustrates the population-level univariate activation estimate associated with each brain parcel (see Figure 3). Asterisks indicate locations of example parcels whose posterior densities are displayed in the right panels. **Right**, Example posterior densities of test–retest correlation values. Dotted vertical lines indicate values of summary statistic reliability estimates. These four regions were chosen as representative examples of the posterior densities within each of the four quadrants in the left panel of this figure (marked by asterisks). *LH ContA IPS 2* is located within rostral aspect of left intraparietal lobule, *RH ContA PFCl 3* within rostral right mid-lateral PFC, *RH LimbicB OFC 3* within right orbitofrontal cortex, and *RH SomMotB S2 8* near right somatomotor and supramarginal gyri.

### 3.5 Multivariate contrasts achieve high individual-level precision by squashing trial-level variability

How do multivariate contrasts improve the certainty of individual-level estimates? We illustrate a means by which this improvement is achieved. Theoretical work has demonstrated that a pivotal quantity for individual difference analyses is the ratio of trial-level variability to individual-level variability (Chen et al., 2021; Figure 1), which we refer to here as the “variability ratio”. When trial-level variability greatly outweighs individual-level variability (a high variability ratio), uncertainty in correlations among individual-level variables is maximized. Conversely, if trial-level variability is reduced, for example, by obtaining measurements less susceptible to such noise, then certainty in correlations will increase. Thus, relative to univariate contrasts, multivariate contrasts may improve certainty by finding a dimension that is less susceptible to trial-level noise than the uniform dimension, to which univariate contrasts are bound (Figure 1).

This hypothesis makes two assumptions. In particular, the uniform dimension should indeed be highly susceptible to trial-level variability. Additionally, the dimension used by the multivariate contrast should be less susceptible to trial-level variability. Are these borne out in our data? Consider an example region in left dorsal prefrontal cortex, which consists of 58 vertices. In this region, BOLD activity can fluctuate across trials in 58 different ways (i.e., along 58 different dimensions). By identifying and ranking these dimensions by the amount of across-trial variability that occurs along them, we can see that a large proportion of the trial-level variability is concentrated in only one or two dimensions (Figure 8, left top, black points and lines). This identification and ranking also lets us assess how the spectrum of trial-level variability relates to the univariate and multivariate contrasts. Each contrast is defined by a weight vector, a particular dimension onto which trial-level activity is projected (see Method). Contrasts that are more susceptible to trial-level variability should be more strongly aligned to high-variability dimensions. Indeed, the univariate weight vector is strongly aligned to the largest dimension of trial-level variability (Figure 8, left top, dark blue). The multivariate weight vector, however, is least aligned to this high-variability dimension, and instead weights other intermediate dimensions more strongly (light blue). These patterns can be summarized within each parcel by computing the total amount of trial-level variability in the direction of each contrast’s weight vector (i.e., by weighting the SD of each dimension by its alignment to univariate or multivariate contrasts, then summing across dimensions). Performing this summarization in every parcel of every subject, we can see that in most of them, univariate contrasts are susceptible to a greater amount of trial-level variability than multivariate contrasts. (Figure 8, left bottom). These findings indicate that a fair amount of the trial-level variability is of low spatial frequency in our data (i.e., relatively uniform across vertices within our brain parcels), and that only multivariate contrasts can exploit this uniformity to find alternative, higher spatial frequency dimensions that are more robust to trial-level variability.

**Figure 8:**
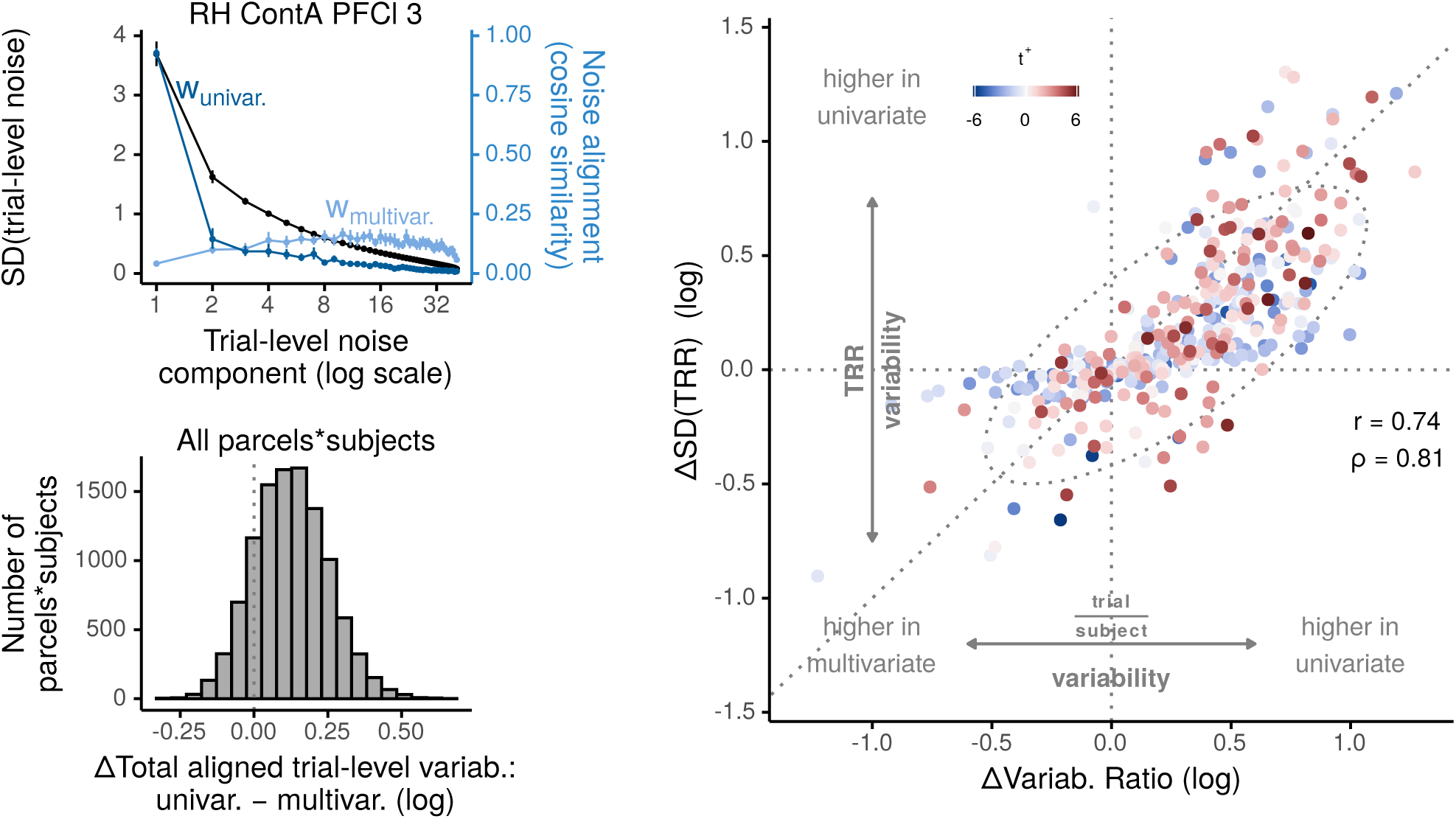
Increased precision through reduced susceptibility to trial-level variability. **Left, top**, Analysis of principal components of trial-level variability and their relation to univariate and multivariate contrasts in an example region. Principal component analysis was used to identify the spectrum of components of trial-level “noise”, that is, the set of dimensions along which this PFC region fluctuates over trials, within each task condition. Each dimension (x axis) corresponds to a particular spatial pattern over vertices within the region, while the black points (left y axis) display how much the region fluctuated along this dimension over trials (in units of standard deviation, or square rooted eigenvalues), on average over subjects. Most variability is concentrated in the first dimension, indicating that trial-level variability was overwhelmingly low dimensional. The blue dots and lines (right y axis) illustrate how these dimensions of trial-level fluctuations are aligned to the univariate (dark blue, **w**_univar._) and multivariate (light blue, **w**_multiv._) weight vectors. The univariate weight vector is primarily aligned to the principal dimension of trial-level variability, whereas the multivariate weight vectors are least aligned to this dimension, and instead more heavily weight intermediate dimensions. For all points, error bars illustrate boostrapped 95% confidence intervals of between-subject variability in means. **Left, bottom**, In most brain parcels and subjects we measured, multivariate weight vectors were susceptible to less total trial-level variability than univariate weight vectors. This susceptibility was estimated separately in each parcel and subject by computing the log ratio of the total amount of trial-level SD in the direction of univariate versus multivariate weight vectors. Positive values indicate univariate weight vectors were aligned to a greater total amount of trial-level variability than multivariate weight vectors. **Right**, Improvements in *trial/subject* variability ratio translated into improvements in certainty of individual-level correlations. The x axis represents log(univariate variability ratio) – log(multivariate variability ratio), while the y axis represents log(univariate SD(TRR)) – log(multivariate SD(TRR)). In other words, positive values indicate that, relative to univariate contrasts, multivariate contrasts either led to a *smaller and thus more favorable* trial/subject variability ratio (x axis), or to a *less uncertain and thus more interpretable* test–retest correlation (y axis). Note that the values and interpretation of the y axis here are equivalent to that of the x axis in Figure 7. For clarity in this figure, however, we relabel this figure’s y axis to match the interpretation of its x axis, wherein higher values of the underlying statistic (SD, trial/subject variability ratio) are less favorable for individual differences analyses.

Having demonstrated the difference in susceptibility to trial-level variability, we then sought to link changes in the trial/subject variability ratio to changes in the certainty of test–retest correlations. Using hierarchical models fitted in each brain parcel, we first computed the trial versus subject variability ratio separately for univariate and multivariate contrasts. Next, to show how multivariate contrasts impact this ratio, we subtracted these ratios between contrast types (after a log transform) to form a change statistic, which is positive when univariate contrasts yielded a higher trial/subject variability ratio, and negative when multivariate contrasts yielded a higher ratio (Figure 8, bottom, x-axis). Following similar logic, we computed a change statistic for the standard deviation of test–retest correlation, such that positive values indicate greater *imprecision* with univariate contrasts, while lower values indicate greater *imprecision* with multivariate contrasts (Figure 8, bottom, y-axis). Most brain regions fall in the all-positive quadrant, indicating that multivariate contrasts made both ratios more favorable for individual difference-focused analyses. Importantly, though, the change in variability ratio was also strongly positively correlated with the change in precision. This relation is consistent with improvements of correlation precision being driven by suppression of trial-level variability in the measures.

## 4 Discussion

We investigated whether a current, widely-used psychological test of cognitive control can yield *reliable* fMRI measures of individual differences over repeated testing sessions. In particular, we focused on characterizing the way in which two contemporary modeling frameworks, Hierarchical Bayesian modeling and MVPA, jointly impact the derived fMRI measures and their associated estimates of test-retest reliability. We found that their combination clearly afforded complementary benefits for estimating individual differences. Hierarchical Bayesian modeling generally led to higher estimates of test-retest reliability than the more traditional “summary-statistic” framework, but in most cases, these estimates were highly uncertain, reflecting strong trial-to-trial variability in the derived measure. This variability, however, was reduced by the application of MVPA, which in turn substantially increased the certainty of the Bayesian estimate of reliability. Therefore, by combining these contemporary modeling frameworks, widely-used task fMRI designs can be used to produce not only highly reliable, but also highly precise measures of individual differences.

### 4.1 Primary conclusions

Our findings caution against sweeping claims of the inadequacy of using task-based fMRI to study individual differences. For example, in one recent study, authors analyzed several extant datasets of common task-based fMRI designs, and reported that the test-retest reliability of univariate activation contrasts in key regions of interest were consistently low (Elliott et al., 2020). These results were interpreted as evidence that “the task-fMRI literature generally has low reliability” and that such task-fMRI measures are “not currently suitable for … individual-differences research” (see also Flournoy et al., 2024). Our findings undermine this conclusion, as we present evidence that one of the most commonly used task designs in fMRI, the color-word Stroop task, can indeed elicit highly reliable fMRI measures.

This discrepancy between findings can be accounted for, perhaps entirely, by our use of hierarchical modeling methods to estimate test-retest reliability, as opposed to their use of summary-statistic methods. When trial-level noise is high — which is invariably the case for fMRI and many behavioral measures of cognition — summary-statistic estimates of reliability are most inaccurate (Figure 1; Chen et al., 2021). In such scenarios, when investigators fail to grapple with these complexities within their approach to data analysis, the risk of faulty and overly pessimistic inferences increases. Our findings join a growing body of work in illustrating the usefulness of hierarchical Bayesian analysis as a principled way of approaching these issues (Chen et al., 2021; Haines et al., 2020; Rouder & Haaf, 2019; Rouder et al., 2023; Snijder et al., 2023). Nevertheless, extending this approach to modeling empirical fMRI data remain a highly infrequent practice (but see, e.g., Chen et al., 2021; Palmeri et al., 2017).

In addition, our findings mark an advance from prior modeling work on individual differences in that we provide evidence for a partial remedy to the “reliability crisis”. Prior work has used hierarchical Bayesian frameworks to pinpoint the primary limiting factor for individual difference studies: the overwhelming influence of trial-level noise (Chen et al., 2021; Rouder et al., 2023). While this insight is invaluable, these studies also demonstrate that hierarchical Bayesian modeling, on its own, does not provide a solution to the psychometric issues gripping this field. In contrast, our findings demonstrate a partial solution to this issue, at least in the case of fMRI, may be provided by MVPA decoding (Kragel et al., 2021). We found that application of MVPA substantially suppressed trial-level noise relative to individual differences, and this led to more certain and interpretable estimates — without compromising the high degree of test-retest reliability. Thus, the use of multivariate rather than univariate contrasts can substantially boost power for detecting individual differences in task-fMRI responses to psychological manipulations.

We have also demonstrated a reason *why* it is the case that MVPA leads to substantially lower trial-level variability than univariate contrasts. Because we used a highly interpretable multivariate framework, linear discriminant analysis, we were able to straightforwardly formulate univariate contrasts as a special case of a multivariate decoder, in which the decoder weights are uniform across vertices (Equation 2). Critically, the only feature that differed between our multivariate and univariate models was the nature of these weights: both models were applied to the same input data, and both implemented a linear scalar projection. The fact that trial-level variability was squashed in our multivariate contrast therefore implies that the MVPA decoder was able to find a dimension (i.e., weights) along which the signal-to-noise ratio was more favorable than the uniform dimension, to which univariate contrasts are bound (see Figure 1 D for a cartoon depiction). Analyses of group-level trends bolster this interpretation, in which we found that the principal component of trial-level variability in many brain regions was strongly aligned to the uniform dimension, whereas the optimal dimension for decoding task conditions was considerably less aligned with the uniform dimension (Figure 8). Whether these improvements result from suppressing noisy vertices (Walther et al., 2016), exploiting noise correlations (Bejjanki et al., 2017; Walther et al., 2016; Zhang et al., 2020), or capturing signal heterogeneity (Davis et al., 2014; Harrison & Tong, 2009; Roth et al., 2018), and the extent to which these features depend on particulars of preprocessing (e.g., smoothing, confound regression, HRF modeling) remain questions for future research. Such questions would be highly tractable to address under the framework we have presented here.

### 4.2 Connections to other work

#### 4.2.1 Joint modeling

Although hierarchical models are, as of yet, used relatively infrequently to model neural timeseries data, there are several notable precedents. Perhaps the most established line of work in this area has been within “joint modeling” a hierarchical modeling framework for integration of neural and behavioral data (Turner et al., 2013, see also Palmeri et al., 2017). Conceptually, the hierarchical modeling aspects of joint modeling are similar to those outlined here, and therefore these approaches likely confer similar benefits for accurate identification of individual differences (e.g., Haines et al., 2023). Additionally, joint modeling approaches can be used to identify multivariate relations among regionally aggregated (i.e., “univariate”) neural timeseries (Turner et al., 2017).

While useful, multivariate joint modeling has not yet been extended to incorporate core features of MVPA decoding: namely, decoders optimized on sensor-level (i.e., voxel-level) input patterns, in a subject-specific manner — in other words, allowing heterogeneity in the decoder weights, across both *subjects* and *sensors within brain region* (see *Future Directions: distributed activation* section in Turner et al., 2019). These features bring several non-trivial differences relative to measures based on univariate contrasts (regional-mean levels of activation, or correlations thereamong). Some are benefits, such as the ability to identify subtle neural encoding of variables within fine-scale, subject-specific, and spatially distributed topographies (Harrison & Tong, 2009; Haxby et al., 2014; Naselaris et al., 2011), increased ability to suppress or enhance distinct components of noise or signal variability (Bejjanki et al., 2017; Hebart & Baker, 2018; Zhang et al., 2020), and a closer bridge to theories of neural population coding and associated statistical methods in computational neuroscience (e.g., Ebitz and Hayden, 2021; Ruff et al., 2018). Others are caveats, such as the curse of dimensionality and unique demands for stratified cross-validation (e.g., Alink et al., 2015). As a result, our work complements prior joint modeling work by characterizing measurement properties of MVPA decoders within a hierarchical model. Indeed, extending joint modeling to incorporate MVPA decoding would likely bring several benefits (see Section 4.4).

#### 4.2.2 Davis et al. (2014)

A relevant prior study compared univariate and multivariate decoding models through simulations from a generative hierarchical model (Davis et al., 2014). Interestingly, these simulations included both trial and individual-level variability, and one conclusion reached was that MVPA may be *less* sensitive to *individual-level* variability than univariate models. This conclusion may initially seem to conflict with ours, in that we found MVPA models can yield more favorable measures for targeting individual differences. Note, however, that this prior study considered a highly restricted case, in which (simulated) individuals only varied in their univariate (uniform) activation to experimental conditions. This assumes that other factors that can strongly influence multivariate decode-ability — for example, the amount of voxel-level variability induced by experimental conditions, or the structure of trial-level variability (e.g., noise correlations) — are not variable across subjects. In real data, however, it seems unlikely that these factors do not differ across subjects. Indeed, even though we did not explicitly optimize the MVPA decoders to enhance individual-level variability (but rather, to enhance condition-level versus trial-level variability within each individual), the resulting output patterns were nevertheless able to strongly individuate participants within our sample.

#### 4.2.3 Marek et al. (2022)

If multivariate decoding indeed boosts power for identifying individual differences in task-fMRI, as we suggest in Section 4.1, then this approach would open several avenues for individual differences research. For instance, a viable research strategy may then be to densely sample a modest number of individuals (i.e., with many trials and sessions), then use MVPA decoding with hierarchical modeling to identify individual differences. Yet at first glance, such a strategy (which requires further validation; see Section 4.4) appears to conflict with prior work claiming that thousands to millions of individuals are necessary to detect individual difference relations between fMRI and behavioral measures (Marek et al., 2022). But these findings are not actually in conflict: rather, they simply reflect different approaches to, and assumptions regarding, the study of individual differences.

The conclusions of Marek et al., 2022 were based primarily on analyses of resting-state and structural MRI, as opposed to task-based fMRI, which was our focus here. In addition, while a small number of task-based fMRI analyses were reported in Marek et al., 2022, the results of those analyses exclusively pertain to what the authors referred to as “brain-wide association studies”, a niche category of individual difference questions, in which all of the covariance between an individual’s fMRI activation contrast from a given task and their behavioral performance on the same task is considered to be noise, and therefore completely statistically discarded (see Extended Data Figure 3 of Marek et al., 2022; Spisak et al., 2023). Such a narrow and unusual definition of individual-difference correlations conflicts with a foundational assumption in psychology: that measures in different tasks are partially generated from a smaller set of latent psychological dimensions, such that dependence on common dimensions drives similarity between measures (Bollen, 2002; DeYoung et al., 2022; Spearman, 1904). As a result, the results from that study have limited to no bearing on the interpretation of results reported here. In fact, as their results show, when this common variance is not statistically discarded, individual-level correlations between task-fMRI and behavioral measures reach moderate effect sizes (*r* s between 0.3 and 0.6) — even despite the use of summary-statistic correlations, which are known to be downwardly biased (Chen et al., 2021), and univariate contrasts, which we have shown here to yield suboptimal precision for individual-level correlations.

#### 4.2.4 Flournoy et al. (2024)

Most recently, Flournoy et al., 2024 examined test–retest reliability of an fMRI contrast associated with perception of fearful facial expressions. This study used a longitudinal dense-sampling design, in which subjects were scanned once monthly for 10 sequential months to contrast the univariate fMRI response to fearful versus neutral expressions. They found that individuals’ fear-related response fluctuated widely from session to session, apparently to the extent that no reliable individual differences were detected in any brain region (generally, ICCs were ∼ 0.1). These reliability estimates, however, were subject to the same interpretational issues as prior work: because they were obtained from trial-aggregated data, they were underestimated (Figure 3, Chen et al., 2021). In addition to the many other substantive differences between that study and ours, (e.g., the nature of the processes targeted by the contrast, use of univariate versus multivariate contrasts), this fact makes these results difficult to compare.

But beyond these differences, Flournoy et al., 2024 illustrates the benefits of a “precision neuroscience” approach, in which a modest number of subjects are intensely characterized in a longitudinal design (e.g., Gordon et al., 2017; Kupers et al., 2024; Naselaris et al., 2021; Poldrack et al., 2015). For instance, although the fear-related response was apparently inconsistent within any given individual, within any given session, an individual’s strength of this response was related to other measures of their current stress level. Thus, slow-timescale affective-state factors (on the order of months) may play an important role for neural responsivity in their task and affective processing more generally.

In contrast to Flournoy et al., 2024, we focused on studying *consistency* across a relatively long timescale (across “test” and “retest” clusters of sessions), and thus did not characterize more transient changes (e.g., across sessions, within cluster). In fact, with our particular cross-validation scheme (see Method), we constrained our multivariate decoders to identify dimensions of neural activity that were stable across sessions. On one hand, this focus may have helped increase the precision of measures at the individual level, by suppressing cross-session variability (as we trained our decoders on data from multiple sessions). Yet on the other, it treats such variability as “noise”, even though it may reflect interesting neural dynamics. For instance, optimal neural coding axes may transform across time due to learning (e.g., Mill and Cole, 2023), representational drift (Driscoll et al., 2017; Ziv et al., 2013), or state-related factors that modulate effective connectivity. With multivariate models, neural coding can be partitioned into different subspaces that are subject to different dynamics. Some subspaces may be highly stable over time, while others may be continuously reorganized. This dichotomy may reflect solutions to computational dilemmas that are highly relevant for cognitive control and task learning (e.g., plasticity versus stability; Driscoll et al., 2022). Future work should seek to understand the factors that drive these transformations, and to link neural variability in each of these subspaces to behavioral dynamics such as learning, or to state-based shifts in functional connectivity (e.g., via Shahshahani et al., 2024). A “precision neuroscience” approach would be well-suited for these goals.

### 4.3 Unexpected results

While our expectations that hierarchical modeling and MVPA leads to benefits in test-retest reliability were borne out in general, there were a minority of brain parcels that did not follow such a pattern. In some cases, there were a group of parcels for which the hierarchically estimated reliability was lower than that estimated through summary statistics. Examining this unorthodox set more closely, we found that they all had near zero or even negative summary statistic reliability (Figure 4). This pattern is consistent with the fact that hierarchical reliability estimates are multiplicatively scaled relative to summary-statistic estimates (Chen et al., 2021). In other cases, MVPA decreased the precision of test-retest reliability compared to univariate analysis (Figure 6). We suspect that this pattern reflects a source of noise to which MVPA is susceptible while univariate analysis is not: changes in the signal or noise topographies across cross-validation splits. Here, for our multivariate models, we cross-validated over data acquisition sessions, which were not only administered on different days, but also involved minor differences in experimental manipulations (see Method 2.7). These factors likely hampered the ability of our decoders to generalize across sessions. As a result, we view our results as providing a relatively pessimistic example of the benefits of multivariate modeling. Future experiments specifically tailored to support this type of analysis, we predict, will yield even more reliable and precise individual difference measures.

### 4.4 Limitations and future directions

Future work would make valuable contributions by exploring several directions that build on these findings.

#### 4.4.1 Linking to behavior

We have demonstrated a framework that can yield highly reliable task-fMRI measures at the individual level. It remains to be seen, however, whether this improved reliability actually translates into improved predictive power for individual differences. Thus, the next clear step will be to use this framework to predict other cognitive or behavioral measures of interest. Finding that MVPA methods are more predictive than univariate methods would provide solid validation of this framework for studying *cognitively relevant* individual differences. This validation would be the strongest if the improved behavioral prediction holds across multiple levels: namely, not only across individuals, but also within individual (i.e., across trials).

Just as we have shown here that a trial-level hierarchical modeling approach is vital for accurate identification of correlations among neural measures, so too will it be for identifying brain–behavior associations, as both types of measures are typically beset by substantial trial-level variability. Such “joint models”, which estimate the (latent) joint distribution of neural and behavioral measures across individuals, afford a natural way to account for variability across these multiple levels (Palmeri et al., 2017). This topic has been been discussed extensively (Turner et al., 2013, 2019), and several recent technological innovations have lowered barriers for implementation (Abril-Pla et al., 2023; Bürkner, 2017; Radev et al., 2023; see also Nunez et al., 2024).

#### 4.4.2 Improved timeseries modeling and hierarchical decoding

We used a simple “selective averaging” method to estimate single-trial activations. This method has many similarities to the popular “least-squares–separate” method (Mumford et al., 2012), and we validated that our single-trial estimates were relatively robust to the particular choice of timeseries model (Supplemental 10). Nevertheless, there are likely more principled and optimized methods that future work could use instead (e.g., Prince et al., 2022). Improvements in estimating single-trial activations would likely translate into substantial improvements in power for identifying individual differences.

Likewise, our spatial models (LDA decoders) and reliability models (hierarchical Bayesian regressions on decoder outputs) were optimized independently from one another. In other words, our decoding method is, in some sense, still a “summary statistic” method, as uncertainty in the decoder weights is not propagated to higher levels of analysis (c.f., Turner et al., 2017). A more accurate approach would be to incorporate the estimation of the MVPA decoder into the hierarchical model itself, amounting to a three-level hierarchical model (trial, voxel, and individual), wherein priors on representational geometry could be placed. Recent developments in Bayesian methods for representational similarity analysis accomplish similar goals (Cai et al., 2019; see also Friston et al., 2019), and have even been extended to encompass estimation of the initial time series model (for discussion of a framework useful for these goals, see Cai et al., 2020).

#### 4.4.3 Decomposing trial-level variability

Another direction worthy of exploration is whether trial-level variability can be usefully decomposed in the service of studying individual differences. Here, we have considered such variability to be “noise”. This consideration was reflected in our use of linear discriminant analysis, which explicitly suppresses trial-level variability. This choice may have obscured dimensions of neural activity that are “behaviorally potent” and linked to cognitively relevant individual differences. For example, if the neural and cognitive processes are subject to cross-trial dynamics, and these dynamics are not adequately captured within the timeseries or decoding model, then the decoder would suppress informative components of neural coding. This suppression may even be expected, given that control and attention processes are non-stationary, fluctuating over trials due to meta-control, distraction, motivation, et cetera (e.g., Saville et al., 2011; Unsworth and Robison, 2017). As such, one may wish to adopt a decoding method that does not suppress such variability (see Kobak et al., 2016 for similar logic) — or, more strongly, to explicitly parameterize and model such variability. Indeed, it is tempting to consider that some of the overwhelming cross-trial variability in neural timeseries such as fMRI (and behavioral measures such as response times) may be explainable in terms of yet-to-be-parameterized neurocognitive processes.

#### 4.4.4 Design innovation

Our results suggest promise for the use of classical experimental designs to study individual differences, but we do not wish to discourage exploration of novel experiments or tasks. Returning to the “drawing board” of task design may bring distinct advantages. Classical designs have been optimized for group-level power, which perhaps occurred at the expense of subject-level power (Hedge et al., 2018). Individual variability may be more strongly elicited by more naturalistic, less constrained, or higher-dimensional designs (e.g., Nastase et al., 2020; Rosenberg and Finn, 2022; Shallice and Burgess, 1991; Sonkusare et al., 2019; Voloh et al., 2023), or by bespoke tasks developed in conjunction with computational cognitive models (Zorowitz & Niv, 2023). But neither of these cases allow one to escape the need to develop sophisticated, theory-driven modeling approaches to make sense of the data. In addition to classical experimental tasks, we predict that novel approaches would be well served by the framework we have illustrated here, which leverages the complementary benefits of MVPA and hierarchical modeling.

## 5 Data and Code Availability

The fMRI data used in the current study comes from the Dual Mechanisms of Cognitive Control Project (Braver et al., 2021). Code and data derivatives sufficient to generate all results and figures reported here are available for download from the following repositories: code doi:10.5281/zenodo.14048490 (archive of https://github.com/mcfreund/trr), data derivatives doi:10.5281/zenodo.14043319. To obtain raw or minimally preprocessed fMRI timeseries data, please contact us so that we can fulfill your requests directly. Raw fMRI timeseries data used in this project are also available for download from the National Institutes of Mental Health Data Archive at https://nda.nih.gov/edit collection.html?id=2970. For detailed descriptions of the task design, see Braver et al., 2021, and of image pre-processing and quality control, see Etzel et al., 2022.

## 6 Author Contributions

Michael C. Freund: *Conceptualization, Formal Analysis, Investigation, Methodology, Software, Visualization, Writing–original draft, Writing–review & editing*. Ruiqi Chen: *Formal Analysis, Methodology, Software, Visualization, Writing–review & editing*. Gang Chen: *Methodology, Writing–review & editing*. Todd S. Braver: *Conceptualization, Funding Acquisition, Resources, Supervision, Writing–review & editing*.

## 9 Acknowledgements

Computations were performed using the facilities of the Washington University Research Computing and Informatics Facility (RCIF). The RCIF has received funding from NIH S10 program grants: 1S10OD025200- 01A1 and 1S10OD030477-01.

We thank all current and former Dual Mechanisms of Cognitive Control team members for their efforts in collecting and curating the dataset on which this study is based. In particular, we thank Kevin Okansen, Leah Newcomer, Erin Gourley, Maria Gehred, Alex Kizhner, Rachel Brough, Al Tay, Catherine Tang, and Anxu Wang for their major efforts in data collection. We thank Carolina Ramirez and Mitch Jeffers for developing the data preprocessing codebase and Nicholas Bloom for developing the experiment software. We thank Joset Etzel for project management and for developing quality control criteria we used to select our sample of participants. In addition, we thank the members of the Cognitive Control and Psychopathology Lab for many fruitful discussions and feedback. M.C.F. thanks Aaron Cochrane for providing helpful comments on a draft of this manuscript. M.C.F. also acknowledges Professor Mike Strube for his foundational instruction in hierarchical modeling, which provided technical insights that contributed to the development of this work.

## 10 Supplementary Material

### 10.1 Validation of the timeseries modeling approach

To estimate single-trial BOLD responses, we used a simple method of selective averaging of responses from TRs at a fixed time-lag from each stimulus onset. Because this method can be inflexible and subject to substantial bias from adjacent trial’s responses, we performed several validation analyses. These analyses confirmed that our design can support selective averaging, that the resulting estimates are straightforwardly interpretable as single-trial evoked responses, and are largely similar to those estimated from other popular estimation procedures.

First, we compare the TRs selected for averaging to aggregate estimates of the deconvolved event-related response contrast between incongruent and congruent trials, as estimated via an FIR model (9 left). This aggregate “Stroop-effect” timecourse (black) was averaged across all vertices within our ROIs, all sessions, and subjects (of a larger sample of N=80). We can see that the selected TRs (grey, vertical dotted lines) correspond to the “peak” of the Stroop-effect response. Additionally, we can see that these TRs correspond well to the peak of a fixed-shape HRF (red), obtained by convolving a 1-second boxcar with AFNI’s *BLOCK* event model, selected based on prior work with this dataset (Freund et al., 2021). Therefore, this window captures the peak of the Stroop contrast well on aggregate, and approximates the time range that would be expected under typical HRF assumptions.

Second, we examine via simple design matrix analysis, how using a convolution approach via regression might compare to use of selective averaging (Figure 9, right). Here, we assume a fixed-shape, 1 s HRF (AFNI’s *BLOCK*). We create a design matrix consisting of an intercept and three separate predicted responses for three adjacent trials, separated in time by 3 intervening TRs (3.6 s), the minimum separation possible within our experiment (Figure 9, right top). We can see that a fair proportion of the middle trial’s predicted response is non-overlapping with those from surrounding trials, and that this lack of overlap is particularly the case within the selected TRs. We see this more directly when computing the pseudoinverse of this design matrix, which illustrates the weights that the regressors assign to each TR (i.e., after accounting for the overlap; Figure 9, right bottom). The TRs that are weighted most strongly are those within the selected window. This suggests that, under the assumptions of this event model, selective averaging of this time window in our design should have a degree of robustness to collinearity. Indeed, when subjects’ design matrices are built using separate *BLOCK* regressors per trial, a majority of trial regressors maintain small variance inflation factors (VIF *<* 3).

**Figure 9:**
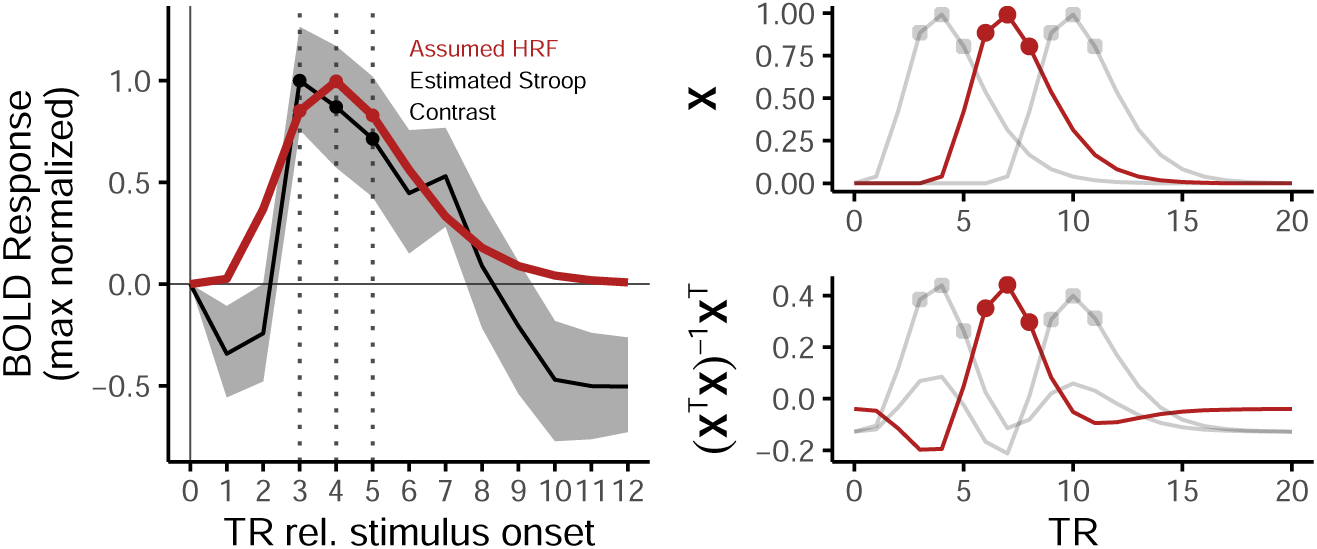
Comparing selected averaging window to deconvolved Stroop contrast and an assumed HRF.

Third, we conduct an empirical sensitivity analysis, in which we compare the estimates obtained from selective averaging to those obtained from two popular approaches for estimating single-trial responses, which we refer to here as “least-squares—separate” (LS-Separate) and “GLM-All with ARMA(1,1).” (see Mumford et al., 2012). These methods differ in several respects from selective averaging and so afford a strong test of sensitivity. LS-Separate is conceptually similar to selective averaging, as both permit correlation among adjacent trial responses, and hence a degree of estimation bias, with the goal of tempering estimation variance that arises from accounting for this collinearity (fitted via AFNI’s *3dLSS*). GLM-All attempts to decorrelate all single-trial responses within a single model, and we fitted this within autoregressive GLMs (AFNI’s *3dREMLfit*, which performs vertex-wise optimization of autoregressive parameters). We compared the selective averaging estimates to each of the other estimates in a variety of ways (Figure 10). In general, selective averaging yields estimates that are similar to both methods, but especially to the popular LS–separate method.

**Figure 10:**
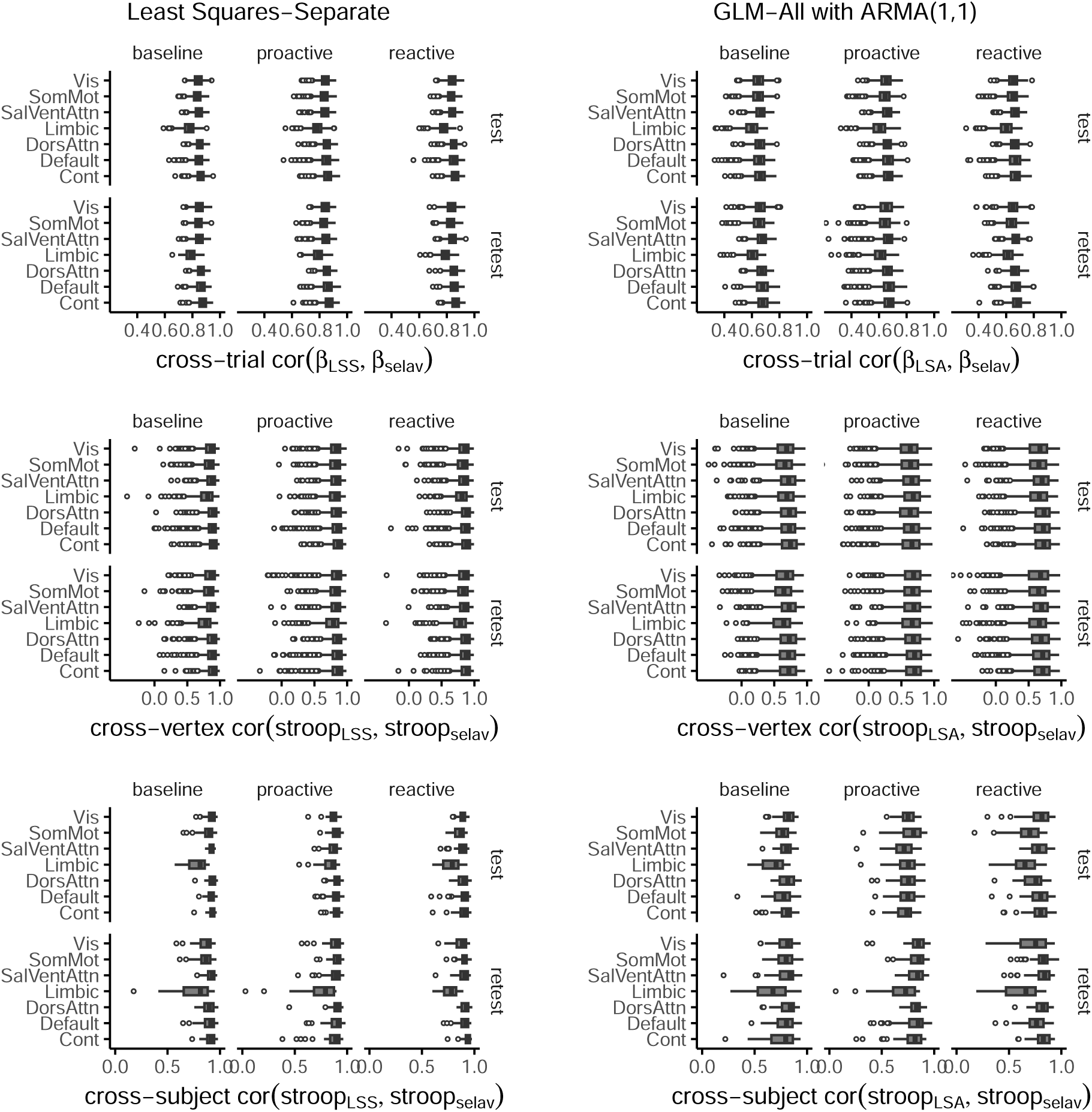
Empirical sensitivity analysis. Estimates from selective averaging were compared to those from two other single-trial timeseries modeling methods: least-squares separate (**left**), and GLM-All with ARMA(1,1) (**right**). All boxplots summarize a distribution of correlations, and are organized by Schaefer-atlas Networks (**y-axes**). **Top**, Within each vertex, parcel, and subject, we estimated the correlation between models in their timeseries of single-trial estimates (i.e., with trials as observations). We then summarized these correlations within each parcel and subject by taking the median over vertices. Boxplots were drawn based on these median correlations (their distribution over parcels*subjects). **Middle**, Within each vertex, parcel, and subject, we estimated the mean Stroop contrast (mean of incongruent minus congruent trials). We then estimated the correlation between models in their spatial pattern of the Stroop contrast estimates (i.e., with vertices as observations). Boxplots were drawn based on these correlations (their distribution over parcels*subjects). (Figure 10, middle). **Bottom**, Within each vertex, parcel, and subject, we estimated the mean Stroop contrast, then took the mean contrast value over vertices. We then estimated the correlation between models in their univariate Stroop contrast estimates (i.e., subjects as observations), per parcel. Boxplots were drawn based on these correlations (their distribution over parcels). 10, bottom).

### 10.2 Model comparison

**Figure 11:**
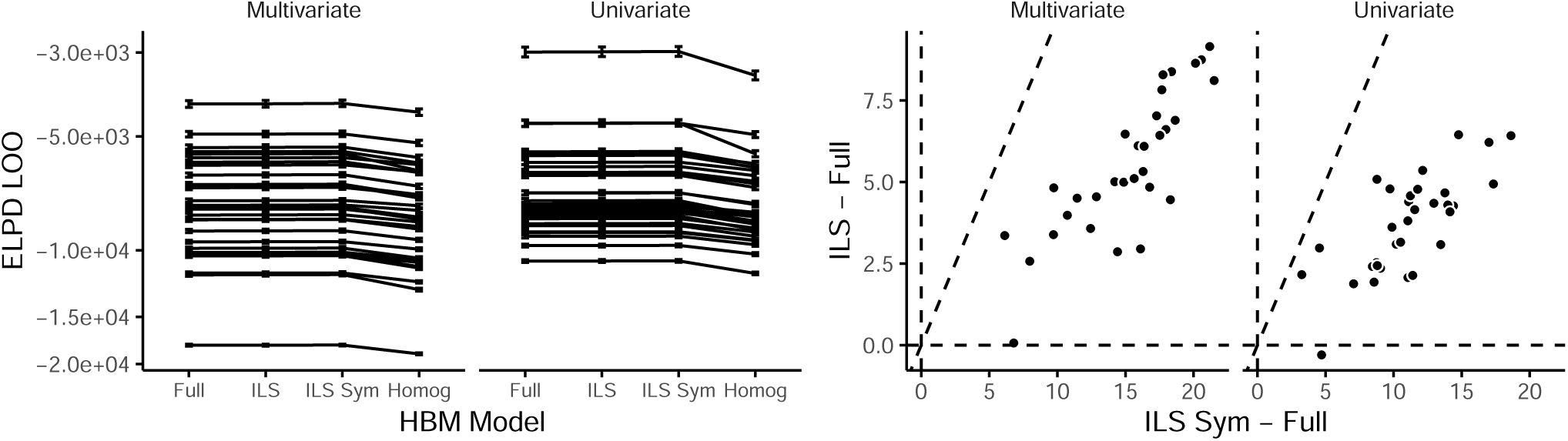
Results of a model comparison. This model comparison was conducted on 32 brain parcels (see Method). Expected-log pointwise predictable density (ELPD) was measured in a leave-one-out manner (LOO). **Left**, Lines connect ELPD LOO estimates (y axis) across different models (x axis) from the same parcel. More positive values indicate a better fit, in terms of better estimated ability of the model to account for out-of-sample datapoints. The pattern of ELPD LOO is highly consistent across parcels. Note that because statistics from the Homog. model were considerably lower (worse) than others, the y-axis spacing is non-linear (inverse hyperbolic sine function). **Right**, Within-parcel contrasts of ELPD LOO. Each point is a parcel. X and y axes illustrate the difference between ELPD LOO for the respective models. Dashed lines illustrate unity line and x and y intercepts. The pattern of ELPD LOO is highly consistent across parcels. On the x-axis, most parcels lie above 0, indicating ILS Sym was preferred over the Full model. On the y-axis, most parcels lie above 0, indicating ILS was preferred over the Full model. Additionally, all parcels lie to the right (underneath) the unity line, indicating ILS Sym was preferred over ILS.

**Figure 12:**
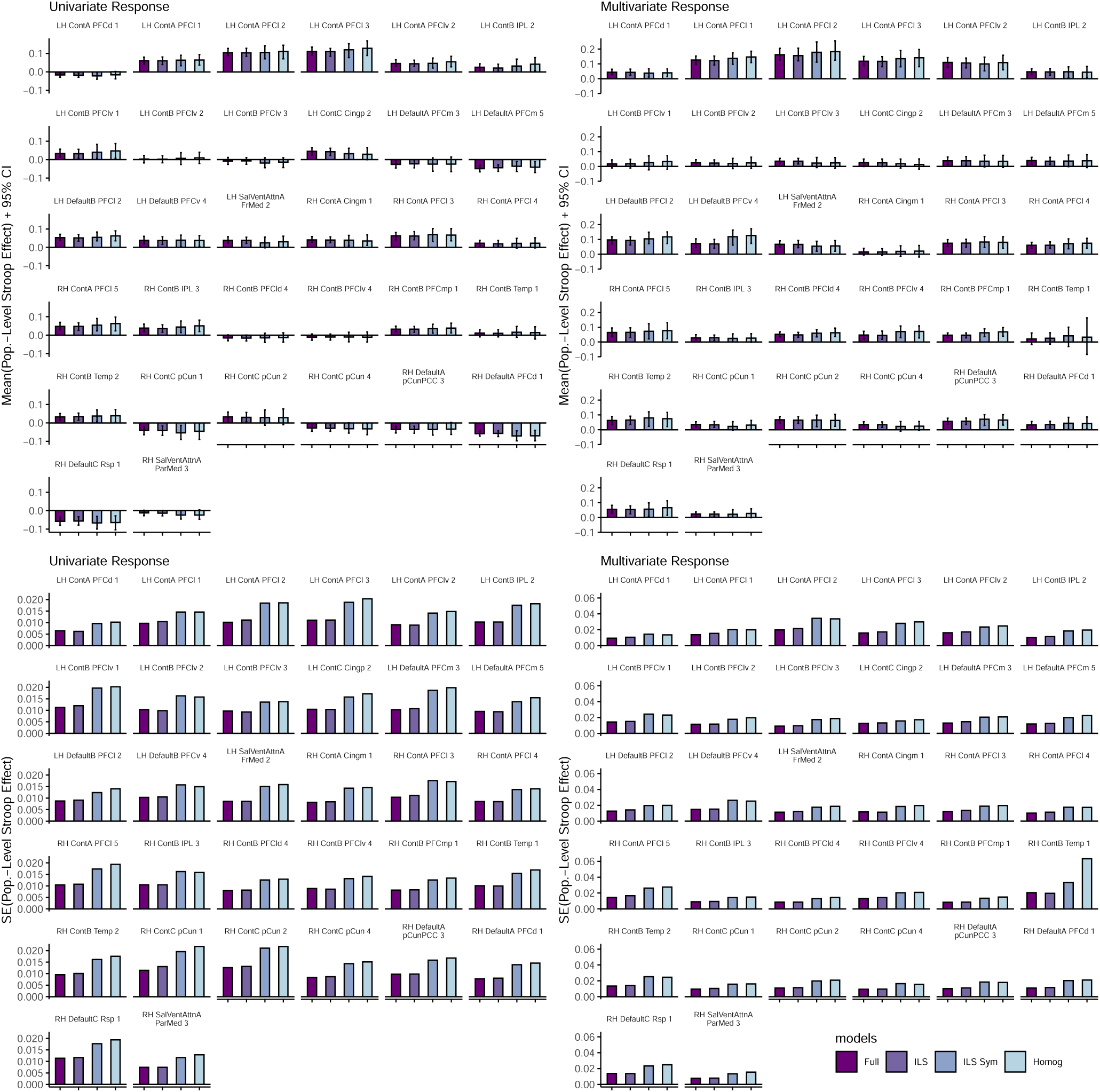
Estimates of population-level Stroop contrasts for all reliability models fitted in 32 brain parcels used for model comparison. **Top**, Bar heights illustrate the mean of the posterior, with errorbars illustrating 95% CI. **Bottom**, Bar heights illustrate the standard error in the mean, measured as the SD of the posterior distribution.

**Figure 13:**
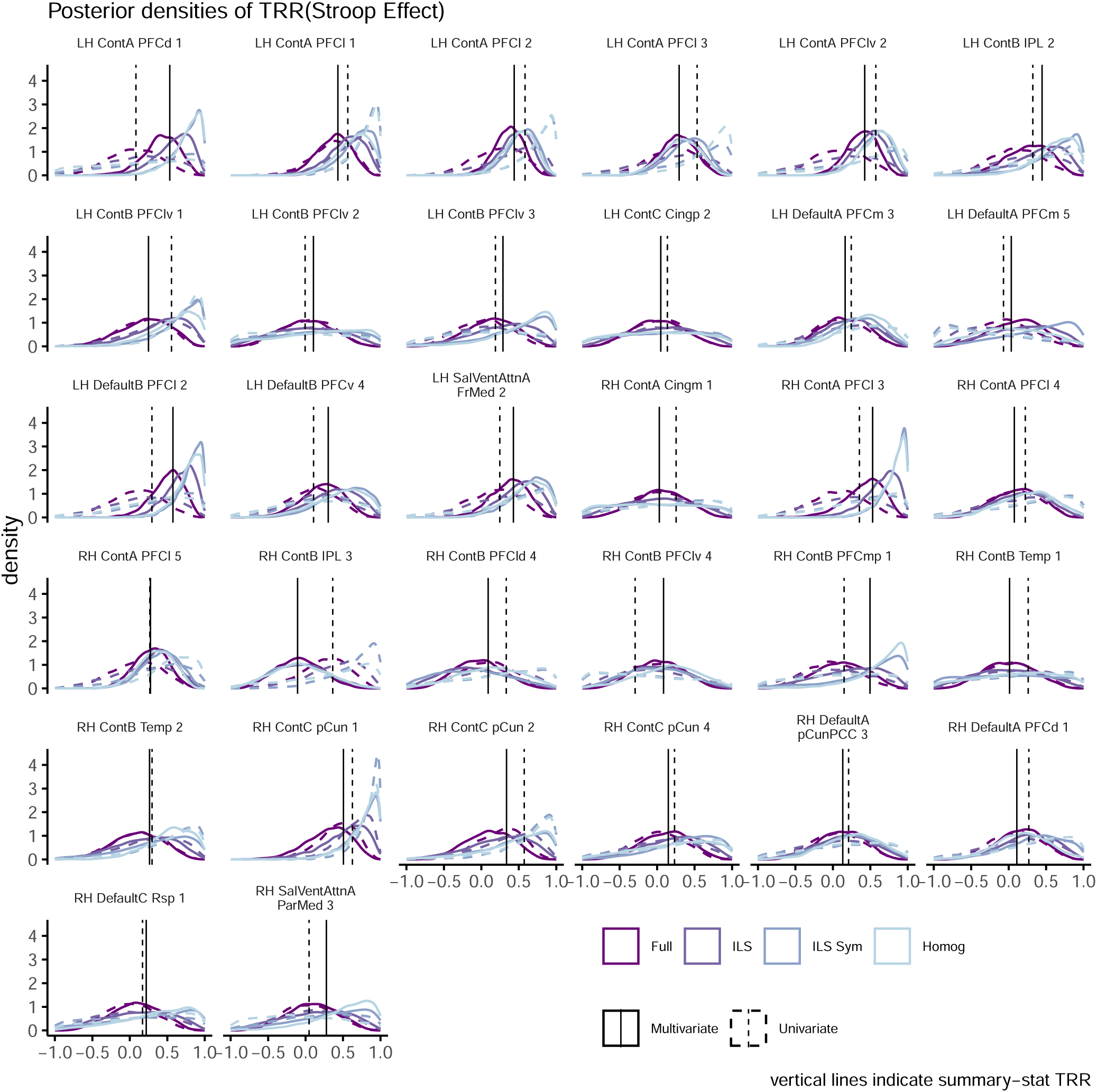
Posterior densities for individual-level test–retest correlations for all reliability models fitted in 32 brain parcels used for model comparison.

**Figure 14:**
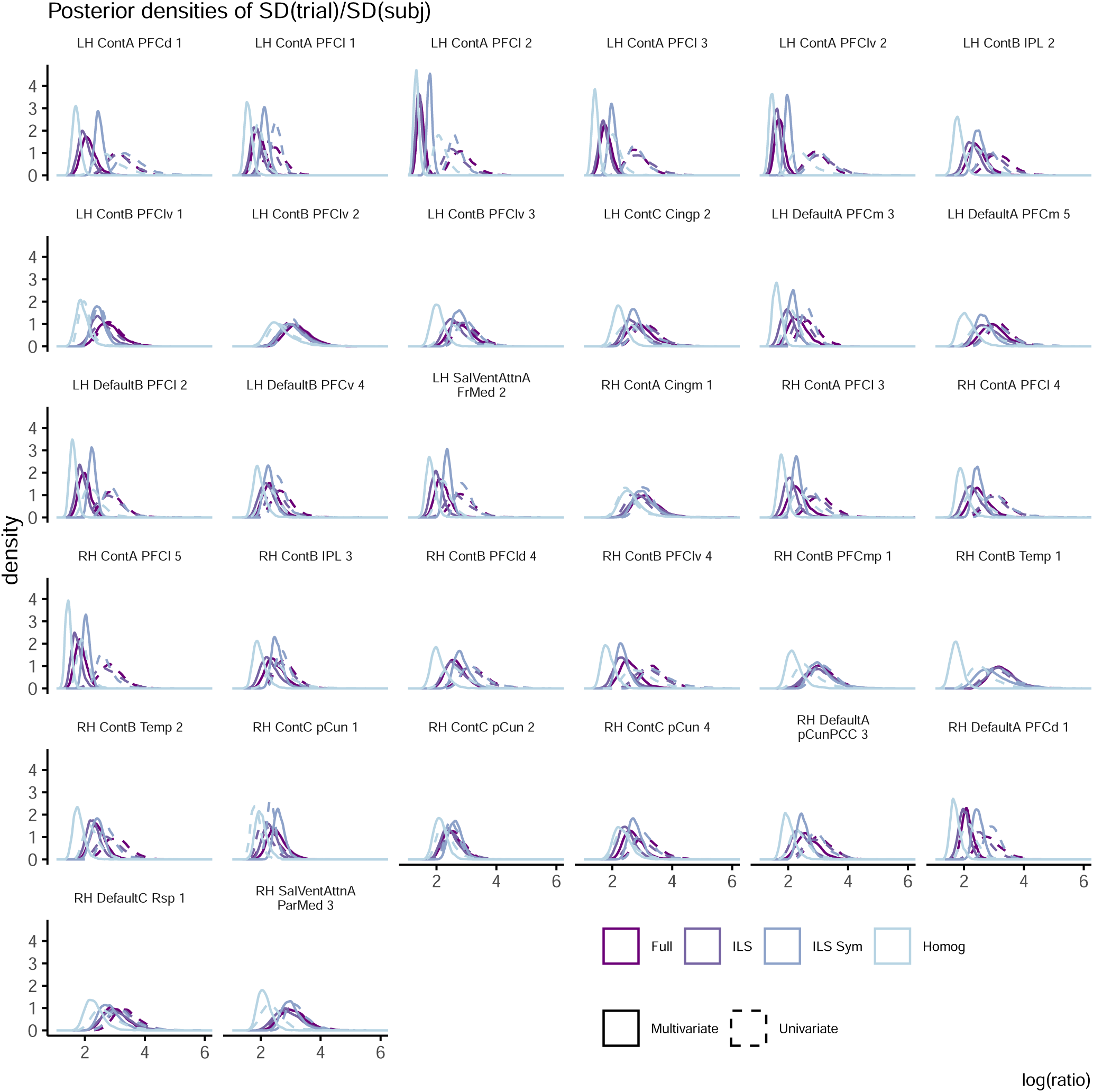
Posterior densities for trial/subject variability ratios for all reliability models fitted in 32 brain parcels used for model comparison.

### 10.3 Supplemental results from selected model (ILS Sym)

**Figure 15:**
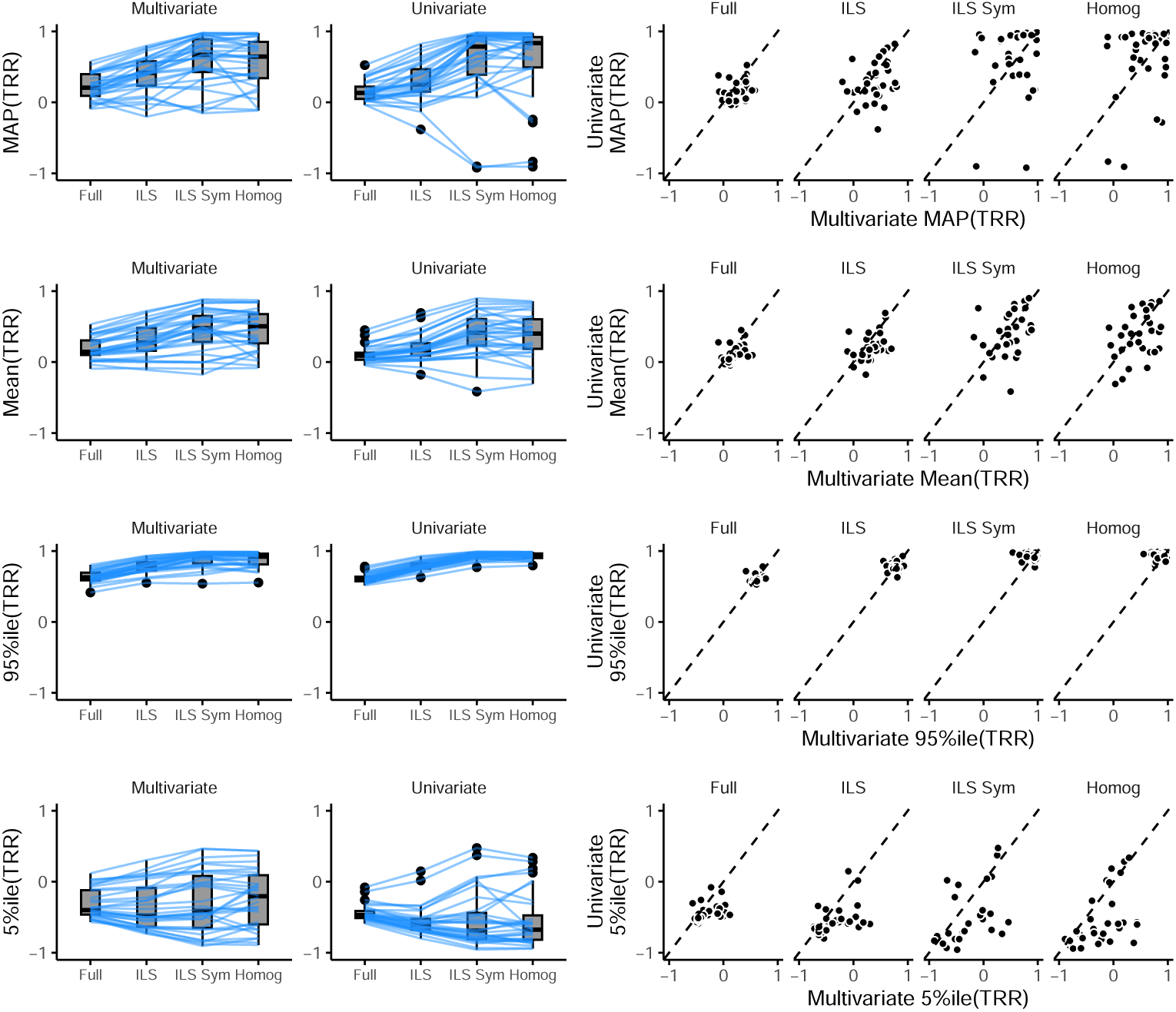
Test–retest reliability statistics from all reliability models fitted in 32 brain parcels used for model comparison. **Left**, On average across 32 *a priori* regions of interest and 5 models we fitted within a model-comparison analysis, reliability estimates tend to be positive, with many *r >* 0.5. Reliability estimates tend to shrink on average when more complex models are fitted (moving right to left on the x axis). As suggested by the model comparison statistics (11), the two most complex models (Full and ILS) are overfitted, while the least complex (Homog.) is underfitted, suggesting results from these models should not bear strongly on inferences. **Right**, Nevertheless, the impact of multivariate decoding relative to univariate contrasts can be seen clearly in the lower-bound estimate (5%-ile) of each model, as most lower-bounds are more positive with multivariate contrasts (x axis) than univariate contrasts (y axis; dashed lines display unity, and most points are below it).

**Figure 16:**
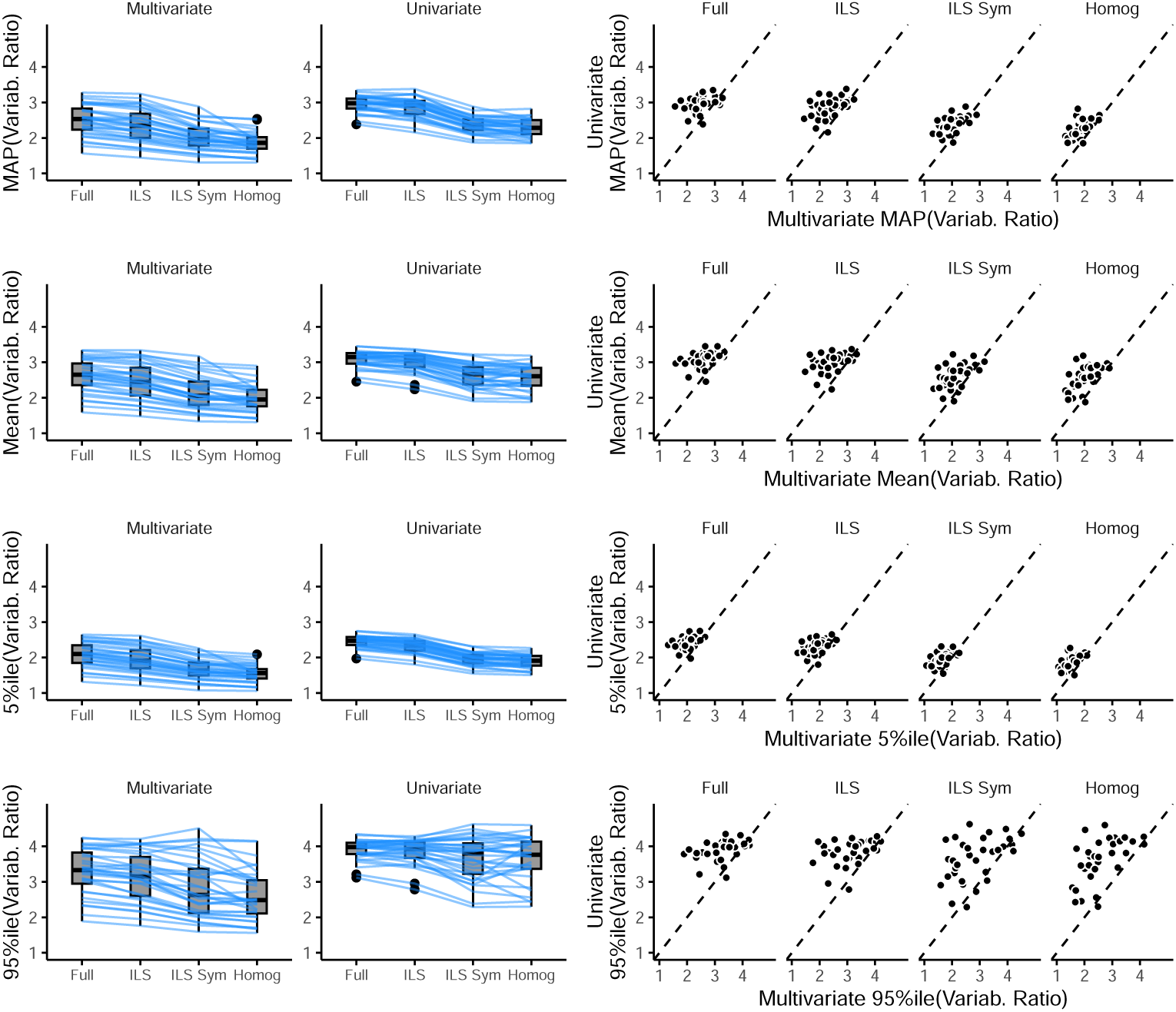
Variability ratio statistics from all reliability models fitted in 32 brain parcels used for model comparison. This figure is analogous to Figure 15. Variability ratio was defined as the log ratio of trial-level variability versus subject-level variability (see Method). Regardless of brain region or reliability model, variability ratios tend to be smaller (i.e., less relative trial-level variability) in multivariate contrasts as opposed to univariate contrasts.

**Figure 17:**
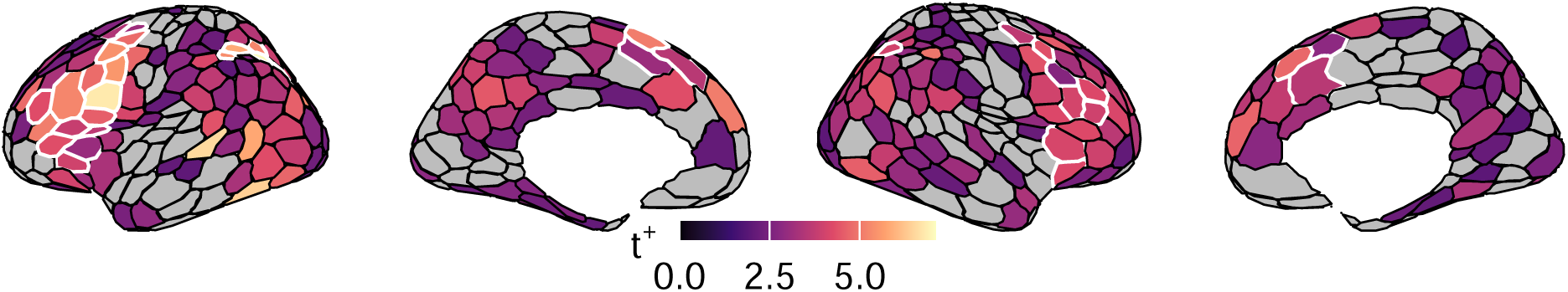
Population-level multivariate contrast. The hue of each parcel depicts the *t*^+^ statistic (see Method or Figure 2), computed on the multivariate Stroop contrast. Only parcels that contained highly discriminable incongruent vs congruent activation patterns are displayed in saturated hues, as defined by a > 0 threshold on the lower-bound of the 95% HDI.

**Figure 18:**
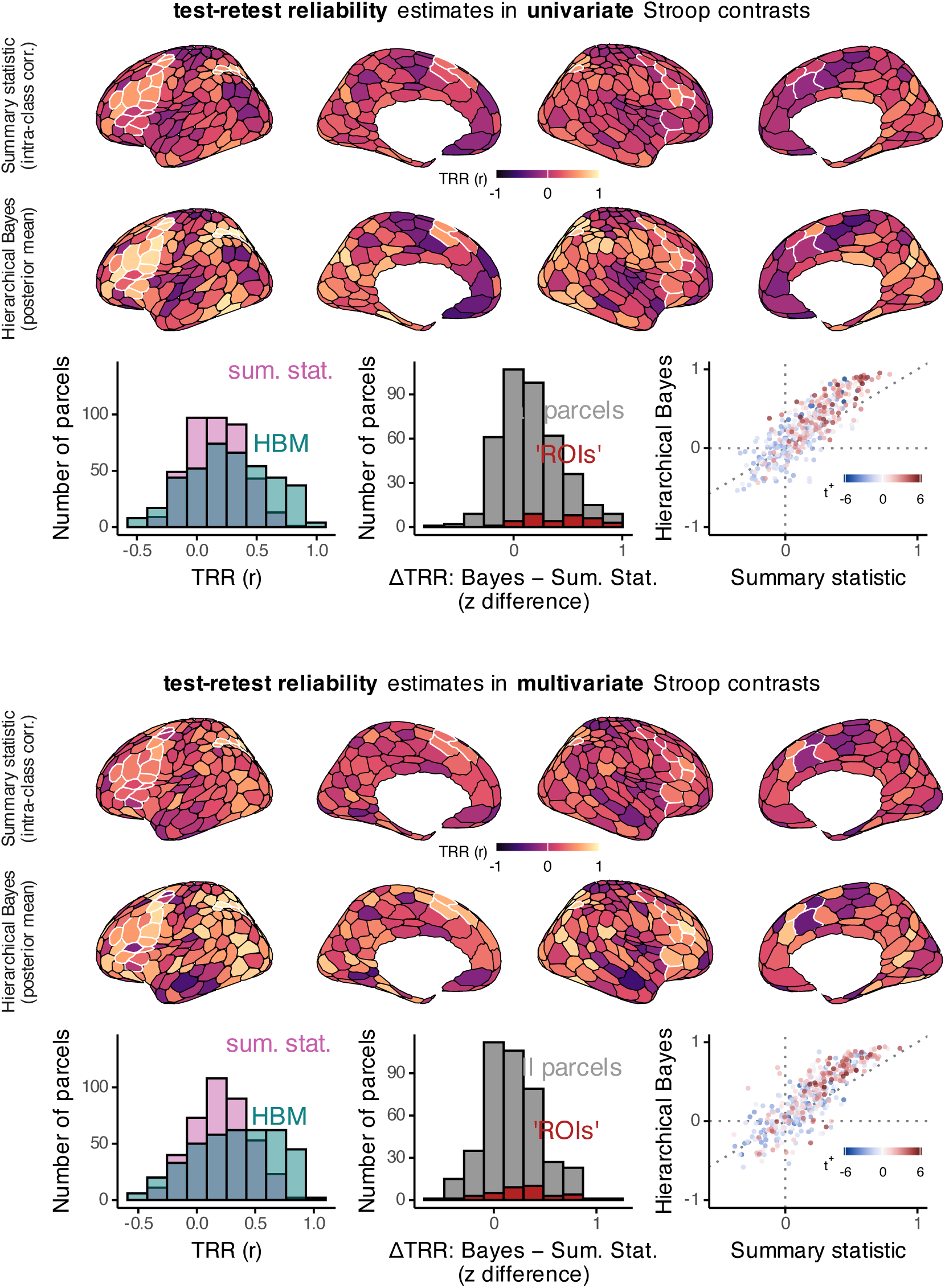
Posterior mean test–retest reliability estimates in univariate (top) multivariate (bottom) Stroop contrasts. All plots are analogous to those in Figures 3 and 4 but using mean rather than MAP.

**Figure 19:**
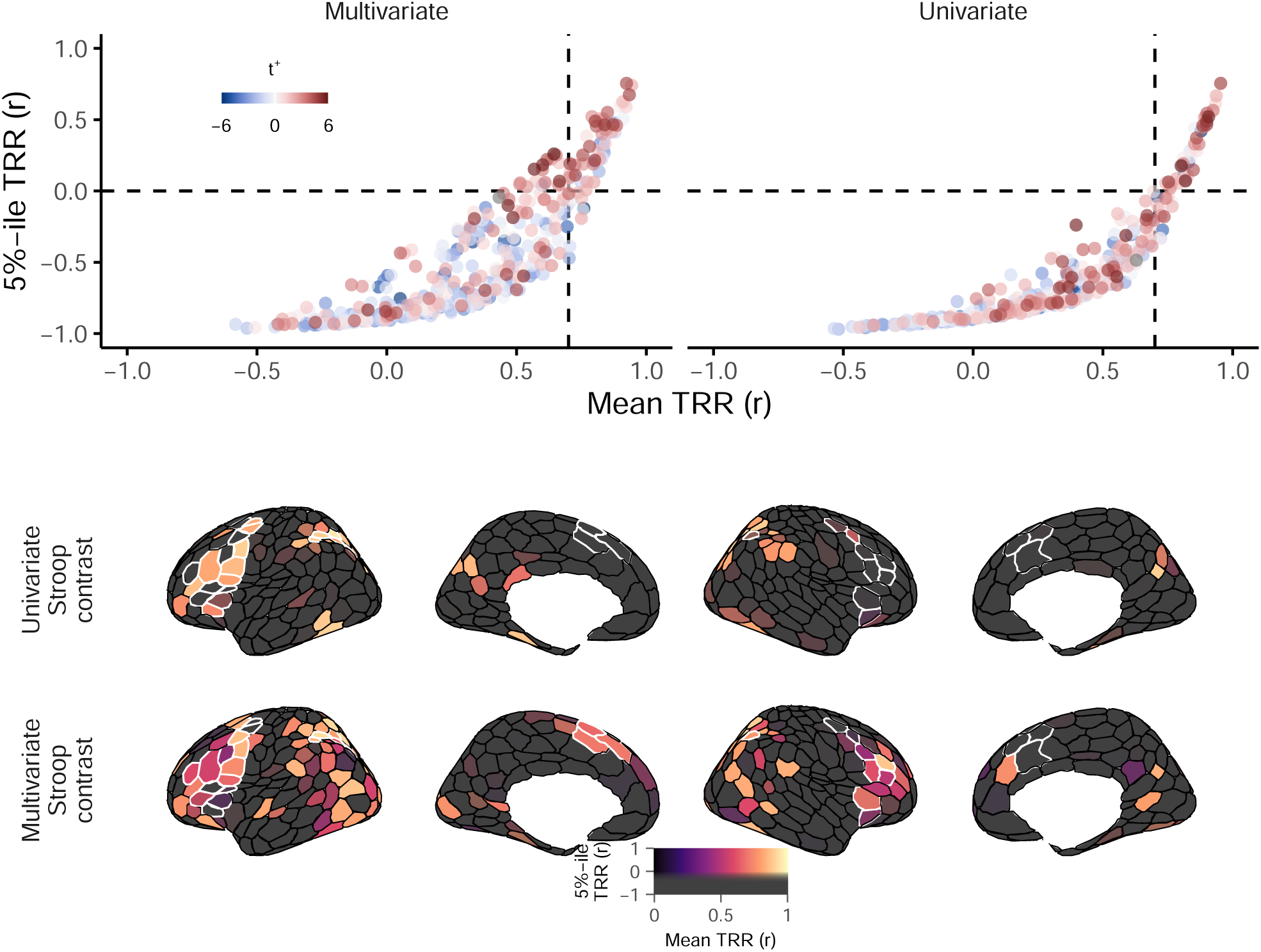
Analogous to Figure 5, but with posterior mean TRR displayed instead.

**Figure 20:**
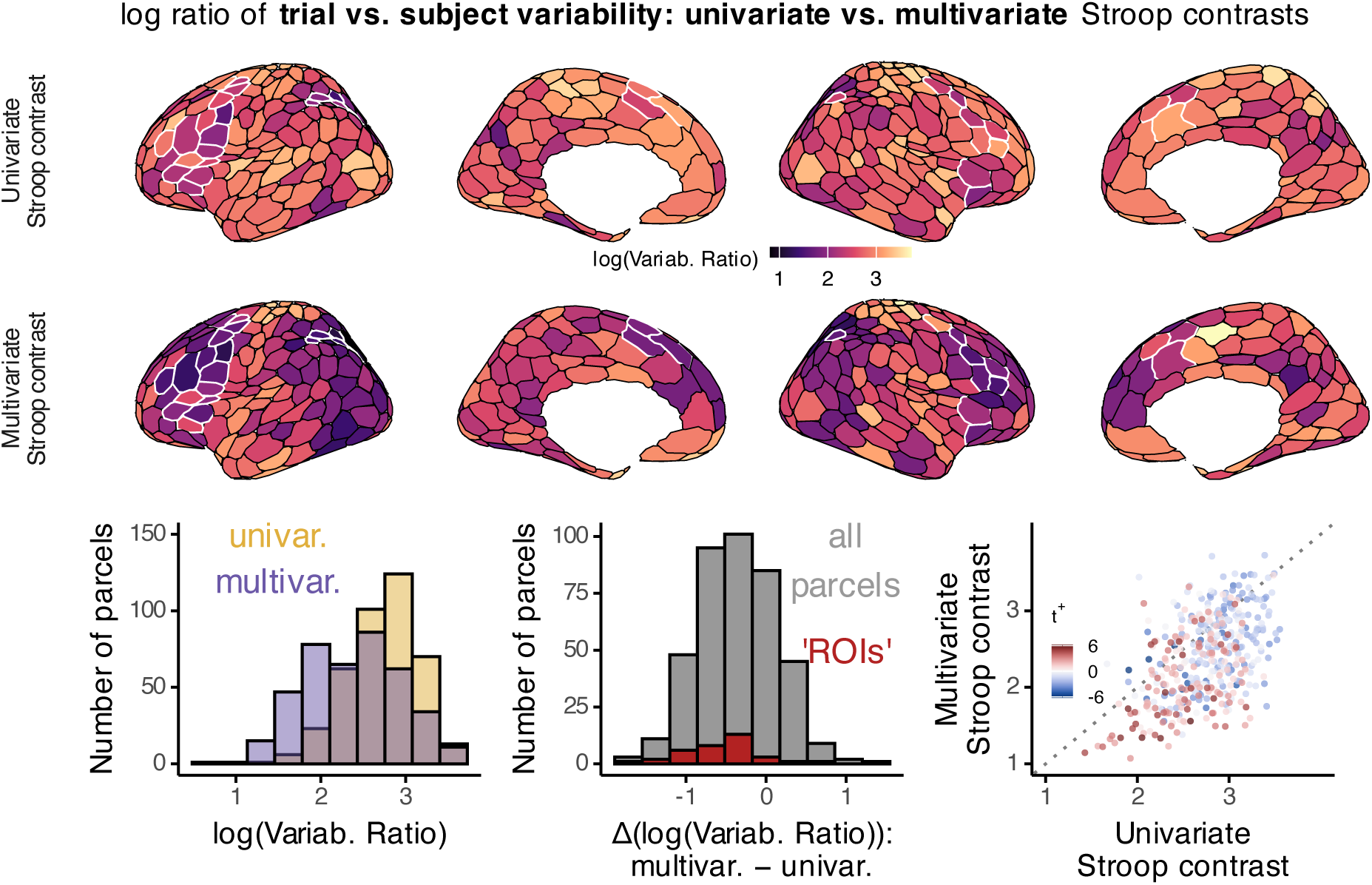
Variability ratio in univariate versus multivariate contrasts.

